# ROS-induced translational regulation—through spatiotemporal differences in codon recognition—is a key driver of brown adipogenesis

**DOI:** 10.1101/2023.12.22.572954

**Authors:** Jun Yu Ip, Indrik Wijaya, Li Ting Lee, Yuhua Lim, Deryn En-Jie Teoh, Cheryl Siew Choo Chan, Liang Cui, Thomas J. Begley, Peter C. Dedon, Huili Guo

**Affiliations:** Institute of Molecular and Cell Biology, Agency for Science, Technology and Research, Singapore; Hilleman Laboratories, Singapore; Genome Institute of Singapore, Agency for Science, Technology and Research, Singapore; NUS Synthetic Biology for Clinical and Technological Innovation, Life Sciences Institute, National University of Singapore, Singapore; Ludwig Institute for Cancer Research, University of Oxford, UK; Department of Biological Sciences, National University of Singapore, Singapore; Antimicrobial Resistance Interdisciplinary Research Group, Singapore-MIT Alliance for Research and Technology, Singapore; Critical Analytics for Manufacturing Personalized-Medicine Interdisciplinary Research Group, Singapore-MIT Alliance for Research and Technology, Singapore; Department of Biological Sciences and The RNA Institute, University at Albany, Albany, NY, USA; Department of Biological Engineering, Massachusetts Institute of Technology, Cambridge, MA, USA; Bia-Echo Asia Centre for Reproductive Longevity and Equality, National University of Singapore, Singapore; Healthy Longevity Translational Research Program, Yong Loo Lin School of Medicine, National University of Singapore, Singapore

## Abstract

The role of translational regulation in brown adipogenesis is relatively unknown. Localized translation of mRNAs encoding mitochondrial components enables swift mitochondrial responses, but whether this occurs during brown adipogenesis, which involves massive mitochondrial biogenesis, has not been explored. Here, we used ribosome profiling and RNA-Seq, coupled with cellular fractionation, to obtain spatiotemporal insights into translational regulation. During brown adipogenesis, a translation bias towards G/C-ending codons is triggered first in the mitochondrial vicinity by reactive oxygen species (ROS), which later spreads to the rest of the cell. This translation bias is induced through ROS modulating the activity of the tRNA modification enzyme, ELP3. Intriguingly, functionally relevant mRNAs, including those encoding ROS scavengers, benefit from this bias; in so doing, ROS-induced translation bias both fuels differentiation and concurrently minimizes oxidative damage. These ROS-induced changes could enable sustained mitochondrial biogenesis during brown adipogenesis, and explain in part, the molecular basis for ROS hormesis.

## Introduction

Brown adipose tissue has been intensively studied in recent years. Unlike white fat which serves as lipid storage sites in the body, brown adipose tissue is specialized to regulate body temperature by burning fat stores to produce heat. This function makes brown fat an attractive target for therapeutics development against metabolic disorders such as diabetes and obesity^1,2^. The discovery that white adipose tissue can undergo “browning” in response to changes in temperature or exercise further highlights the importance of this area of research—if white adipose tissue can be induced to become beige fat and serve the same fat-burning purposes, then they too, can be harnessed to treat metabolic disorders^1,2^. Large-scale mitochondrial biogenesis is a hallmark of brown adipogenesis and browning—understanding how this can be effected would thus lead to therapeutic insights.

Of the >1,000 components needed to make mitochondria, only 13 are encoded within the mitochondrial DNA^3^. The remaining components are expressed from the nucleus and eventually assembled into functional mitochondria. The coordination involved in this process is traditionally studied at the level of transcription. Brown fat differentiation is driven by a cascade of events starting from a network of transcription regulators that is anchored by PRDM16, C/EBP-β, PPARγ, and PGC1-α^1^. The ensuing mitochondrial biogenesis results in cells that are densely packed with mitochondria^4^, giving the cells a brown appearance. It is estimated that mitochondria produce ∼90% of cellular ROS as by-products of electron transport^5^. The ramping up of mitochondrial biogenesis during brown adipogenesis thus inevitably raises intracellular ROS levels. While excessive ROS are highly damaging to cells, there has been increasing evidence that ROS play physiological roles in many biological contexts^6^. Mitochondria-produced ROS are required for activating PPARγ-dependent transcription during white adipocyte differentiation^7^. In mature brown adipocytes, mitochondrial ROS cause oxidative sulfenylation of a cysteine residue in UCP1, which in turn induces thermogenesis^8^. Whether changes in ROS have a direct involvement in brown adipogenesis remains to be explored.

Even less is understood on the role of translational regulation in brown adipogenesis. A previous ribosome profiling study found that brown adipogenesis is driven primarily by changes in RNA abundance^9^. However, whether there were more nuanced aspects of translational regulation was not explored. Proximity-specific ribosome profiling carried out in *S. cerevisiae* has demonstrated that 87% of mRNAs encoding mitochondrial components are localized to the mitochondrial surface^10^. Such localization was later also demonstrated in HEK293T cells using proximity-specific ribosome labeling^11^ and the APEX-seq technique^12^, which does not rely on ribosome association. In *Drosophila,* some of these mitochondrial component-encoding mRNAs are localized by PINK1, a mitochondrial outer membrane kinase. Upon localization to the outer membrane, these originally translationally repressed mRNAs are relieved of repression by PINK1 and Parkin to facilitate protein synthesis^13^. Localized translation enables proteins to be synthesized at the sites where they are most needed, thus facilitating swift responses^14^. Whether the massive mitochondrial biogenesis that occurs during brown adipogenesis employs similar mechanisms to express mRNAs encoding mitochondrial components is not known.

Here, we use ribosome profiling and RNA-Seq, in conjunction with cellular fractionation, to explore how translational regulation might be harnessed to drive brown adipogenesis. In particular, we explore the role of ROS in inducing this process through influencing the activity of ELP3, a tRNA modification enzyme.

## Results

To understand the molecular changes involved in brown adipogenesis, we made use of an established mouse brown preadipocyte cell line as a model system^15^. Over a six-day differentiation time course, the preadipocytes differentiated into mature brown fat cells with abundant mitochondria. To obtain spatiotemporal information on translational regulation during brown fat differentiation, we employed ribosome profiling and RNA-Seq, together with cellular fractionation, to examine samples harvested daily (Figure 1A). Ribosome profiling and RNA-Seq were applied in parallel to clarified lysates from the bulk cytoplasm (without nucleus), the crude mitochondria fraction, or the cytosolic fraction (without mitochondria). This level of granularity yielded much more detailed information compared to typical ribosome profiling studies. In the bulk cytoplasm, translational changes can be observed from Day 4 onwards, with mitochondrial components tending to be upregulated, which is expected as cells ramp up mitochondrial biogenesis (Figure 1B). However, analysis of data from crude mitochondria fractions reveal that translational changes were actually taking place from Day 2 onwards (Figure 1C). These changes were masked in the bulk cytoplasm because the same mRNAs in the cytosolic fraction do not exhibit much translational change (Figure 1D). Translational changes thus appear to start off first in the mitochondrial vicinity before becoming visible at the whole-cell level. In contrast, at the mRNA level, changes can be observed in the early stages of differentiation, regardless of the cellular compartment examined (Figure S1A–C). As expected, mitochondrial components are also upregulated at the RNA level, as the cells massively upregulate mitochondria once differentiation is induced.

**Figure 1.**
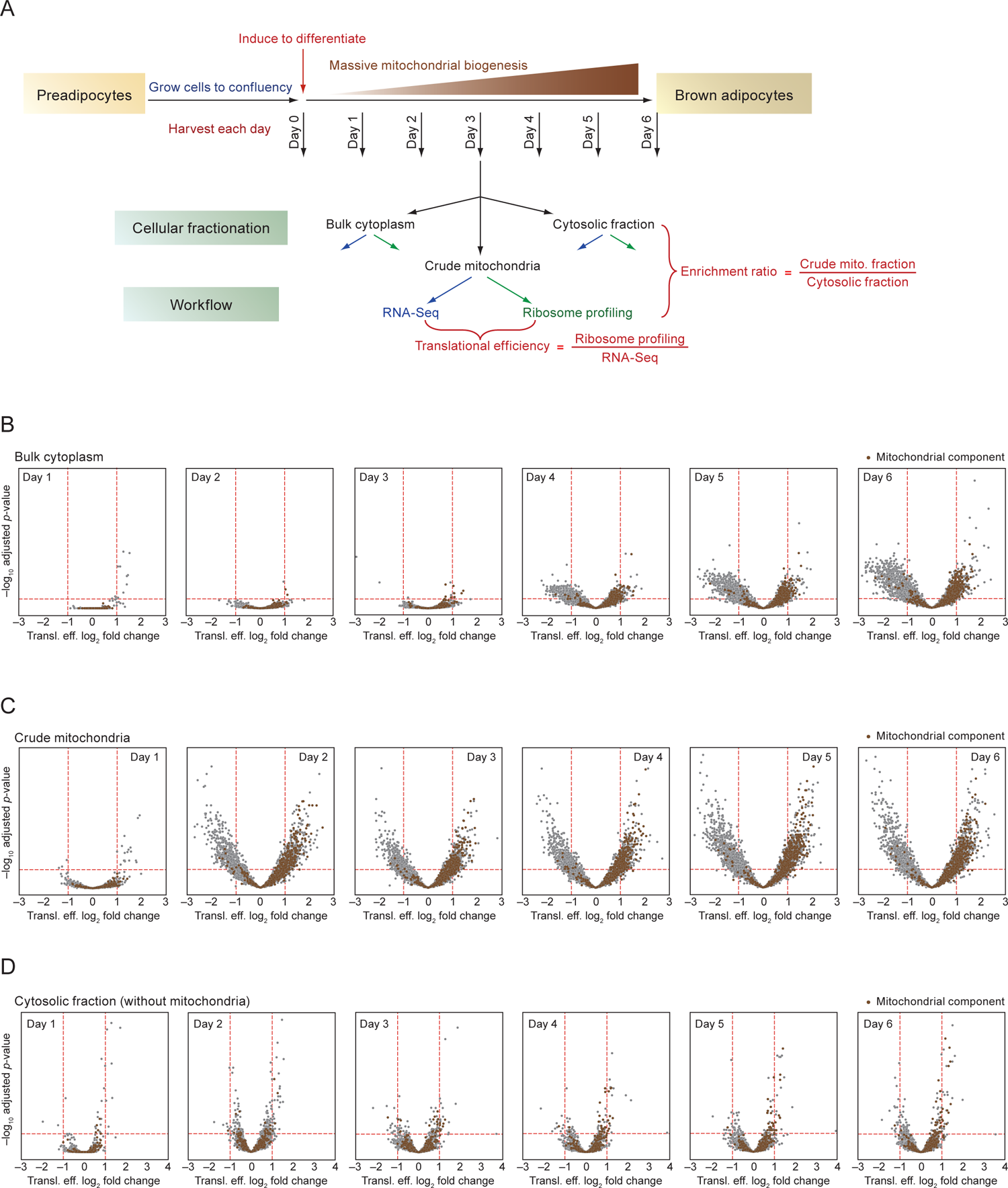
Translational regulation takes place in the vicinity of mitochondria early on in brown fat differentiation. **(A)** Experimental approach taken to probe spatiotemporal changes in RNA and translation levels during brown adipogenesis. Preadipocytes differentiate into mature brown fat cells over six days. On each day, harvested cells were fractionated into crude mitochondria and cytosolic fractions. Parallel RNA-Seq and ribosome profiling were performed on clarified lysates from the bulk cytoplasm (without nucleus), the crude mitochondria fraction, and the cytosolic fraction. **(B)** Volcano plots showing translational efficiency (transl. eff.) changes in the bulk cytoplasm for each day (relative to Day 0) as brown adipogenesis progresses. Genes encoding mitochondrial components are highlighted in brown. **(C)** Translational efficiency changes in the crude mitochondria fraction. Otherwise, as in **(B)**. **(D)** Translational efficiency changes in the cytosolic fraction. Otherwise, as in **(B)**.

To determine whether there is any relationship between the observed translational changes and the codons found in the mRNAs present, we applied the following methodology: we plotted the translational efficiency fold change of each mRNA against the frequency of a particular codon in the mRNA (Figure 2A). This yielded a correlation coefficient, which we term “codon translation coefficient (CTC)”, that was then plotted in a consolidated bar plot to aid visualization for all 61 coding codons (Figure 2B). Analyzed this way, a hierarchy of the relationship between codon frequencies and translational changes emerged. In the crude mitochondria fraction, the more G/C-ending codons (GC3 codons) that an mRNA has, the more translationally upregulated it tends to be (Figure 2C). In the cytosolic fraction, the opposite trend was seen early on in differentiation, i.e. mRNAs with more codons that end in A/U (AU3 codons) were favored (Figure 2D). However, midway through differentiation (Day 4), there appears to be a switchover, after which mRNAs with more GC3 codons became favored, resembling the scenario seen in the crude mitochondria samples. These data suggest the presence of a signal causing such a translation bias, which starts off near the vicinity of mitochondria and later spreads to the rest of the cell, such that by Day 5, the translation bias in the cytosolic fraction then resembles that seen in the crude mitochondria fraction.

**Figure 2.**
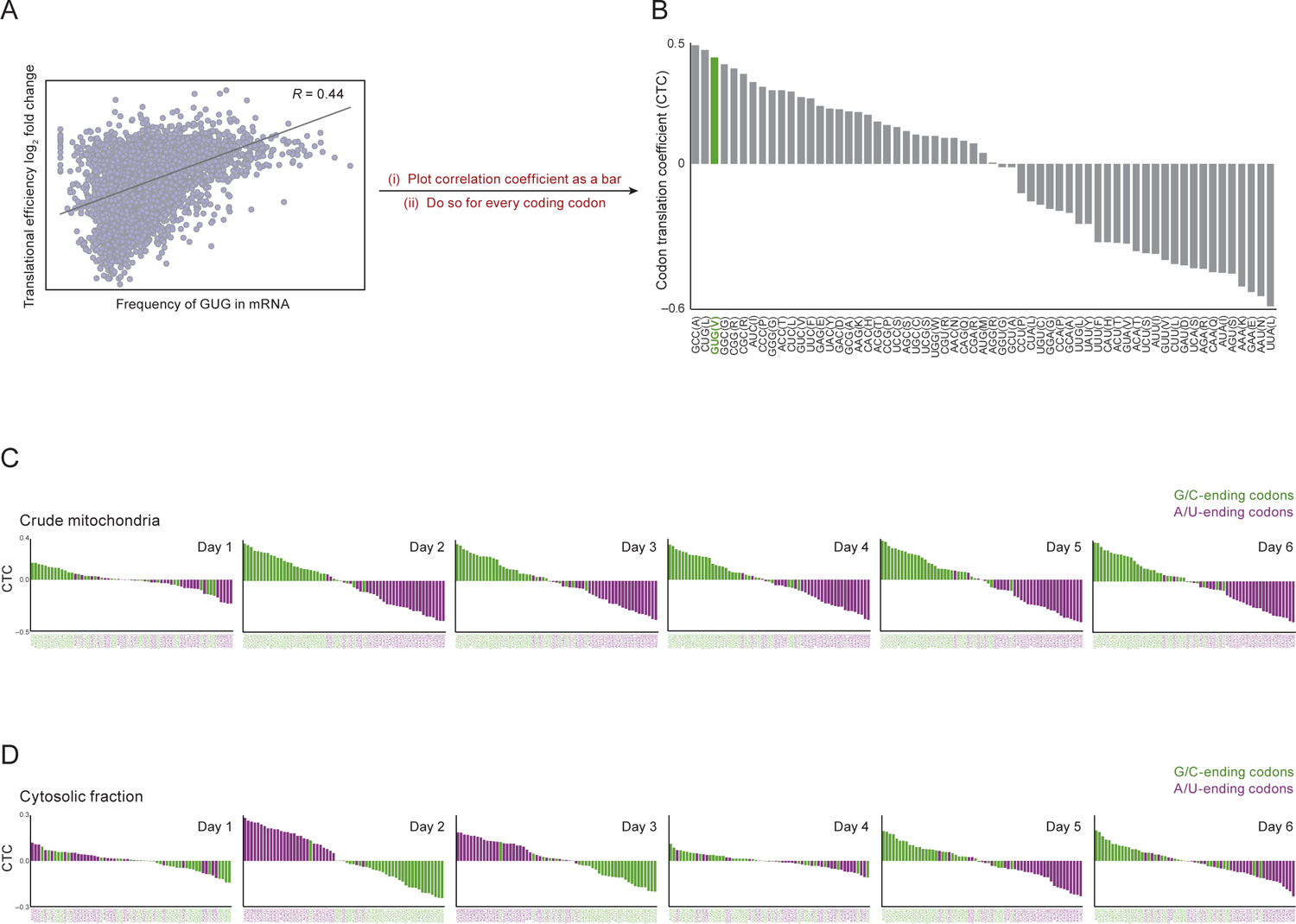
A translation bias towards G/C-ending codons is observed during brown adipogenesis that first originates in the vicinity of mitochondria. **(A)** Scatter plot showing translational efficiency changes and the codon frequency of GUG in mRNAs. The Pearson correlation coefficient is indicated. **(B)** Bar plot of correlation translation coefficients (CTCs) for all coding codons. For each codon, a scatter plot was made plotting log_2_ fold change in translational efficiency against codon frequency as in **(A)**. The correlation coefficient obtained from the scatter plot was then plotted as a bar to aid visualization for all 61 coding codons. Relative fold changes from Day 6 (bulk cytoplasm) were used. The correlation coefficient of GUG codon from **(A)** is highlighted in green. **(C)** Bar plots of CTCs for translational efficiency changes in the crude mitochondria fraction from six days of differentiation (relative to Day 0). G/C-ending codons are colored green. A/U-ending codons are colored purple. **(D)** Bar plots of CTCs for translational efficiency changes in the cytosolic fraction from six days of differentiation. Otherwise, as in **(C)**.

This translation bias is not limited to any single amino acid but is an across-the-board bias towards GC3 codons. A translation bias towards G/C in the third codon nucleotide position (the wobble base) suggests that there might be changes in the cellular pool of tRNAs. We thus examined tRNA abundance in the crude mitochondria and cytosolic fractions by quantitative RT-PCR but did not detect coordinate upregulation of GC3 codon-recognizing tRNAs that could explain the observed translation bias (Figures S2A and S2B). Because the translation bias was first seen in the crude mitochondria fraction, before spreading to the cytosolic fraction (Figures 2C and 2D), we also sought to determine if tRNAs recognizing GC3 codons were upregulated first in the crude mitochondria fraction, but did not detect this (Figure S2C). As such, changes in tRNA abundance could not explain the observed translation bias. tRNAs are known to be extensively modified with nearly 50 different chemical structures of the epitranscriptome, and modifications to the anticodon stem loop can alter the way codons are recognized^16^. We thus used mass spectrometry to measure tRNA modifications in order to identify modified ribonucleotides that could lead to the observed translation bias. An initial survey of tRNA modifications was inconclusive (Figures S3A and S3B). We then searched the literature for tRNA modification enzymes whose activity could explain such across-the-board bias towards mRNAs with more GC3 codons. This led us to ELP3, a key component of the Elongator complex, which is known to modify tRNAs at position 34, the position that recognizes the wobble base. Within the Elongator complex, ELP3 is the effector enzyme that adds the initial modification, a 5-carboxymethyl group (cm^5^), to uridine at position 34 of tRNAs (Figure 3A). Subsequent enzymes add other modifications, leading eventually to four final modifications affecting 11 tRNA isoacceptors, potentially modulating the translation of 22 possible codons^17^ (Figure S3C). Four out of these 11 tRNAs—tRNA-Lys(UUU), tRNA-Gln(UUG), tRNA-Glu(UUC) and tRNA-Arg(UCU)—have been reported to change their codon recognition preference to favor A-ending codons upon modification of uridine-34^18–20^ (Figure 3A). Although such behavior has yet to be demonstrated for the remaining seven tRNAs, we reasoned that if the remaining seven are similarly affected, then since ELP3 can modify 11 tRNAs, modulating ELP3 alone during differentiation would produce an apparent translation bias towards G-ending codons. Therefore, we continued to pursue ELP3 as a prime candidate that could explain our observations. Because we observed preferential translation towards G-ending codons during brown adipogenesis, we predicted that there would be a reduction in ELP3-mediated modifications as differentiation progresses (Figure 3A). Indeed, this was observed when we measured ELP3-mediated tRNA modifications during differentiation (Figure 3B). Intriguingly, ELP3-mediated modification levels were more reduced in tRNAs from the crude mitochondria fraction than those from the cytosolic fraction, which is consistent with our observation that the translation bias towards G-ending codons is observed first, and to a larger extent, in the crude mitochondria fraction (Figure 2).

**Figure 3.**
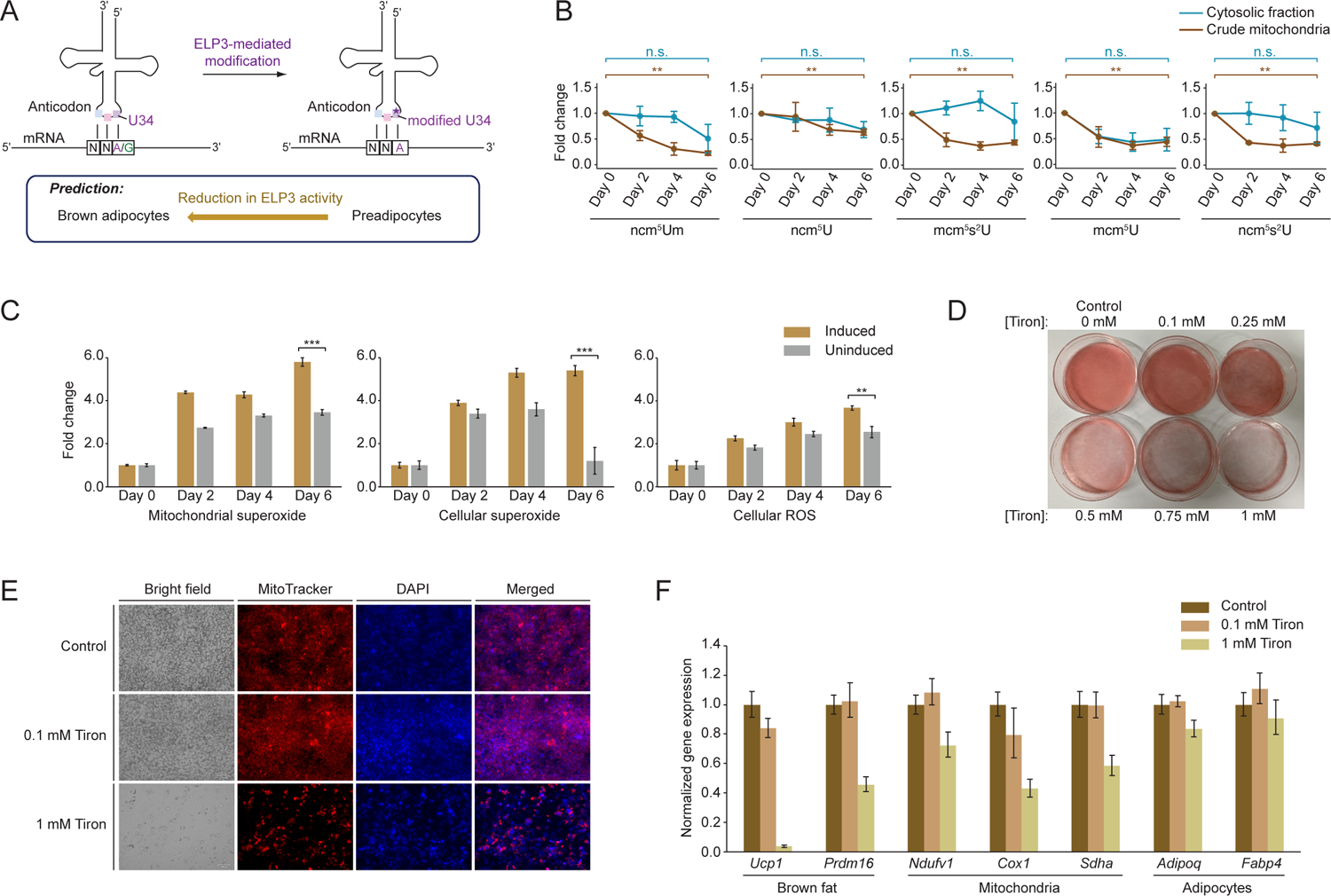
Reduction in ELP3-mediated tRNA modifications is consistent with the production of a translation bias towards G-ending codons during brown adipogenesis, which requires a certain level of ROS. **(A)** Schematic showing modification of tRNAs by ELP3. When U34 is modified, the tRNA preferentially recognizes A at the wobble base (see text). In the context of brown adipogenesis, ELP3 activity is expected to decrease as differentiation progresses, which will lead to an increase in the proportion of unmodified tRNAs and in turn, an apparent translation bias towards G-ending codons. **(B)** Mass spectrometry measurements of ELP3-mediated tRNA modifications. Data shown is the average of three biological replicates, with the corresponding standard deviations (error bars). The first four modifications shown are the four known final modifications added to ELP3-affected tRNAs (Figure S3C). The fifth modification (ncm^5^s^2^U) has been reported to be a potential intermediate in the formation of mcm^5^s^2^U, and is thus also included as an ELP3-mediated modification^59^. The Student’s t-test was used to compare modification levels between Day 6 and Day 0 samples [**: *p* < 0.01, n.s. = not significant]. **(C)** ROS measurements in induced or uninduced preadipocytes. Mitochondrial superoxide, cellular superoxide or cellular ROS (including superoxide) levels were measured in uninduced and induced cells. Data shown is the average of six technical replicates. Error bars represent standard error of the mean. The data shown is representative of two biological replicates. The Student’s t-test was used to determine if ROS levels at Day 6 are higher in induced cells [***: *p* < 0.001, **: *p* < 0.01]. **(D)** Oil Red O staining of cells that have been induced to differentiate in the presence of different concentrations of Tiron. Oil Red O stains lipids red. **(E)** MitoTracker staining of live cells that have been induced to differentiate in the presence of different concentrations of Tiron. Cells were counter-stained with 4’,6-diamidino-2-phenylindole (DAPI). **(F)** Quantitative RT-PCR measurements of known expression markers for brown fat, mitochondria and adipocytes. RNA was isolated from cells treated with the indicated concentrations of Tiron. *Tbp* was used as the internal control. Data shown are fold differences calculated using the ΔΔC_T_ method, derived from three biological replicates, with the standard deviations represented as error bars.

These observations suggest that the signal that leads to a translation bias starts off first in the vicinity of mitochondria and then later spreads to the rest of the cell. One possibility for this signal is ROS produced from the mitochondria. ROS levels are expected to increase during differentiation due to increased mitochondrial biogenesis. Indeed, upon differentiation induction, ROS levels increased more than when cells were uninduced (Figure 3C). Intriguingly, ELP3 is an iron-sulfur (Fe-S) cluster-containing enzyme^21^. Its activity is thus expected to be modulated by ROS levels, because ROS can change the oxidation state of the iron in the Fe-S cluster, leading to enzyme inactivation^22^. This would give us a plausible working hypothesis to explain the translation bias: as differentiation progresses, increased mitochondrial biogenesis raises ROS levels, which in turn reduces the activity of ELP3 enzyme; this increases the proportion of unmodified tRNAs present, leading to a translation bias towards G-ending codons. Should this ROS-induced translation bias be necessary for brown adipogenesis, it would predict that having a certain level of ROS is necessary for proper differentiation. Indeed, when an ROS scavenger, Tiron, is added to preadipocytes, they failed to differentiate normally and accumulated less lipids and mitochondria (Figures 3D and 3E). Expression levels of mRNA markers for brown fat, mitochondria and adipocytes were also reduced when Tiron is added (Figure 3F). These results indicate that a certain level of ROS is required for brown adipogenesis.

Up to this point, we had inferred a translation bias through analyzing sequencing data—i.e. as differentiation progresses, we see more ribosome-protected footprints coming from mRNAs with more GC3 codons. Such an increase could be observed due to genuine increased translation of mRNAs with more GC3 codons. However, there is also a possibility that the increase could be due to slowed or stalled ribosomes over such mRNAs. To distinguish between the two scenarios, we made use of luciferase reporters of different codon compositions. Because the wobble base can be changed without changing the encoded amino acid, it is possible to design reporter constructs that encode the same luciferase protein yet are made up of codons that are differentially regulated by changes in ELP3 activity. Hence, we designed different variants of firefly luciferase as such: ELP3-affected codons that do not end in G were replaced with the corresponding G-ending codons that encode the same amino acid, to produce a G-maximized luciferase; conversely, ELP3-affected codons that do not end in A were replaced with the corresponding A-ending codons to produce an A-maximized luciferase (Figure 4A). These modified firefly luciferase constructs were then transfected into the cells as they were differentiating, and luciferase activity was assessed 24 h after transfection. Consistent with the translation bias that we observed from sequencing, the G-maximized luciferase was translated better than the A-maximized luciferase as differentiation progresses (Figure 4B). Our working hypothesis predicts that this translation bias is induced by ROS. As such, we should observe a decrease in the translation bias towards mRNAs with more G-ending codons (G-bias) if we were to deplete ROS from the system. To test this prediction, we added Tiron to the cells before and during differentiation. With Tiron addition, the G-bias did not simply decrease, but the translation bias completely shifted towards favoring the firefly luciferase reporter with more A-ending codons (Figure 4C). These results indicate that the G-bias is induced by ROS, through modulating ELP3 activity.

**Figure 4.**
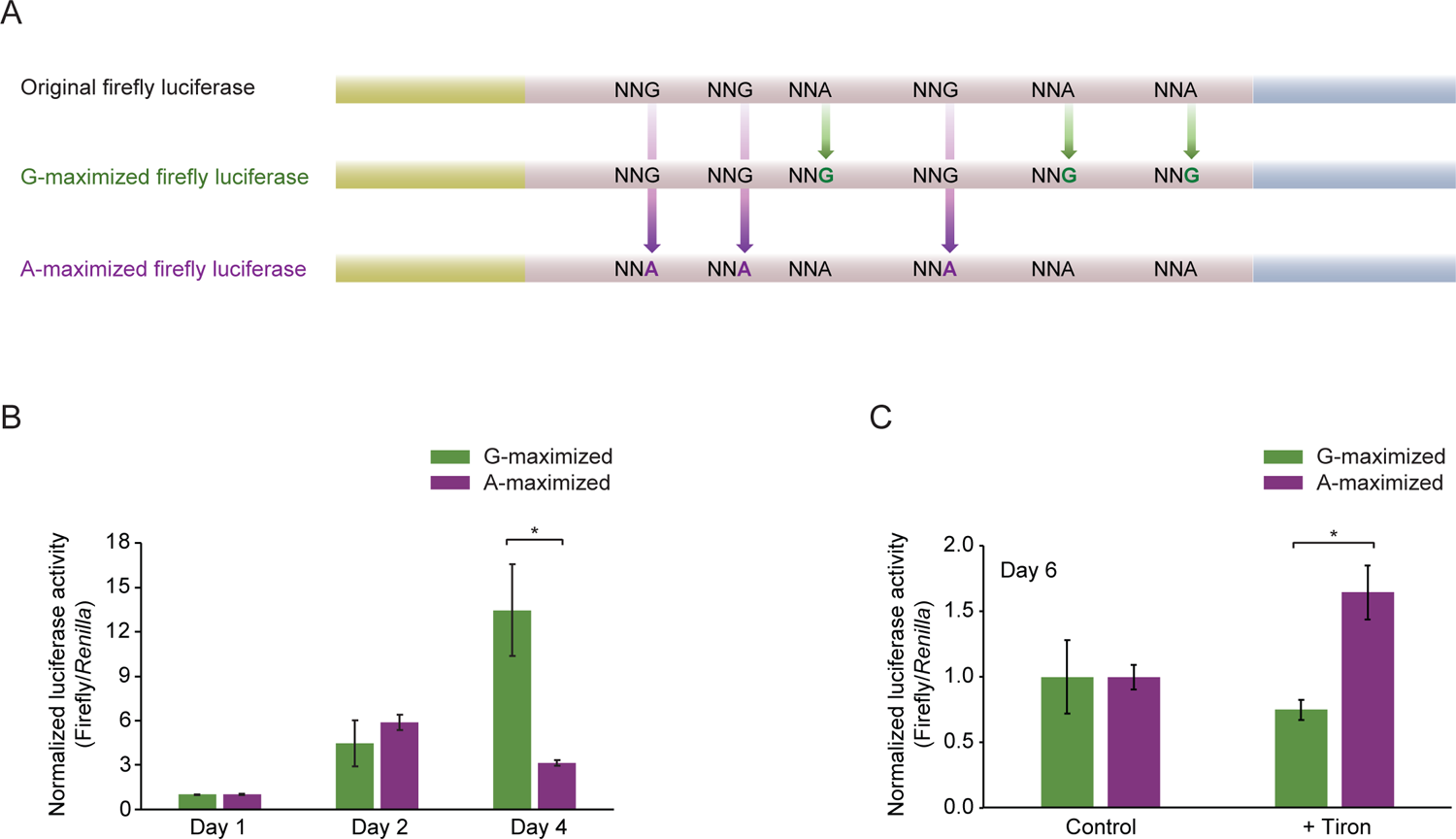
ROS-induced changes in ELP3 activity are responsible for the translation bias towards G-ending codons during brown adipogenesis. **(A)** Firefly luciferase constructs used in reporter assays. ELP3-affected codons were mutated to obtain a G-maximized luciferase or an A-maximized luciferase. Changing the wobble base does not change the amino acid. **(B)** Dual luciferase assays for reporter constructs transfected into cells during brown adipogenesis. The indicated firefly luciferase construct was transfected into differentiating cells 24 h before cells were harvested for dual luciferase assays. *Renilla* luciferase was co-transfected as a control. Data shown is the average of three biological replicates. Error bars represent standard error of the mean. The Student’s t-test was used to compare luciferase activity between the G-maximized reporter construct and the A-maximized one [*: *p* < 0.05]. **(C)** Dual luciferase assays for reporter constructs transfected into differentiating cells, under normal, or ROS-depleted conditions. In the case of ROS depletion, cells were kept in media containing 1 mM Tiron prior to, and during, differentiation. On Day 5, the indicated firefly luciferase was transfected, along with *Renilla* luciferase as a control. Cells were harvested 24 h later (Day 6) for dual luciferase assays. Otherwise, as in **(B)**.

A key question to address is how this ROS-induced translational regulation might drive brown adipogenesis. This phenomenon would contribute to brown adipogenesis if mRNAs encoding brown fat differentiation pathway components or mitochondrial components benefit from the translation bias. We thus probed whether GC3 codons are over-used in certain functional groups of genes. To this end, we calculated the extent to which codons are over- or under-represented in each open reading frame (ORF) in the mouse genome, using a Z-score matrix^23,24^. >60% of genes had at least one codon that was significantly over- or under-represented [codon Z-score > 2 or < –2, respectively] (Figure 5A). Within this group of genes, we demarcated four clusters with different extents of GC3- or AU3-codon biases. We then asked whether certain functional groups of genes are enriched in the cluster exhibiting strong GC3-bias (Cluster 2). Concurrently, we ranked all the genes according to their aggregate Z-scores. This allows us to place different groups of genes along a one-dimensional hierarchy that depicts the degree to which genes over-use GC3 codons (Figure 5B). Brown fat differentiation pathway components are enriched in Cluster 2 and rank highly in GC3 codon over-usage. For mitochondrial components, certain functional groups, e.g. mitochondrial import proteins and respiratory complexes, over-use GC3 codons, while other groups such as enzymes involved in fatty acid β-oxidation and the tricarboxylic acid (TCA) cycle tend to be under-represented in Cluster 2, and rank low on the hierarchy (Figures 5B and S4A). Interestingly, the gene groups that rank lower tend to encode components found in the mitochondrial matrix, while those that rank higher tend to encode mitochondrial membrane components.

**Figure 5.**
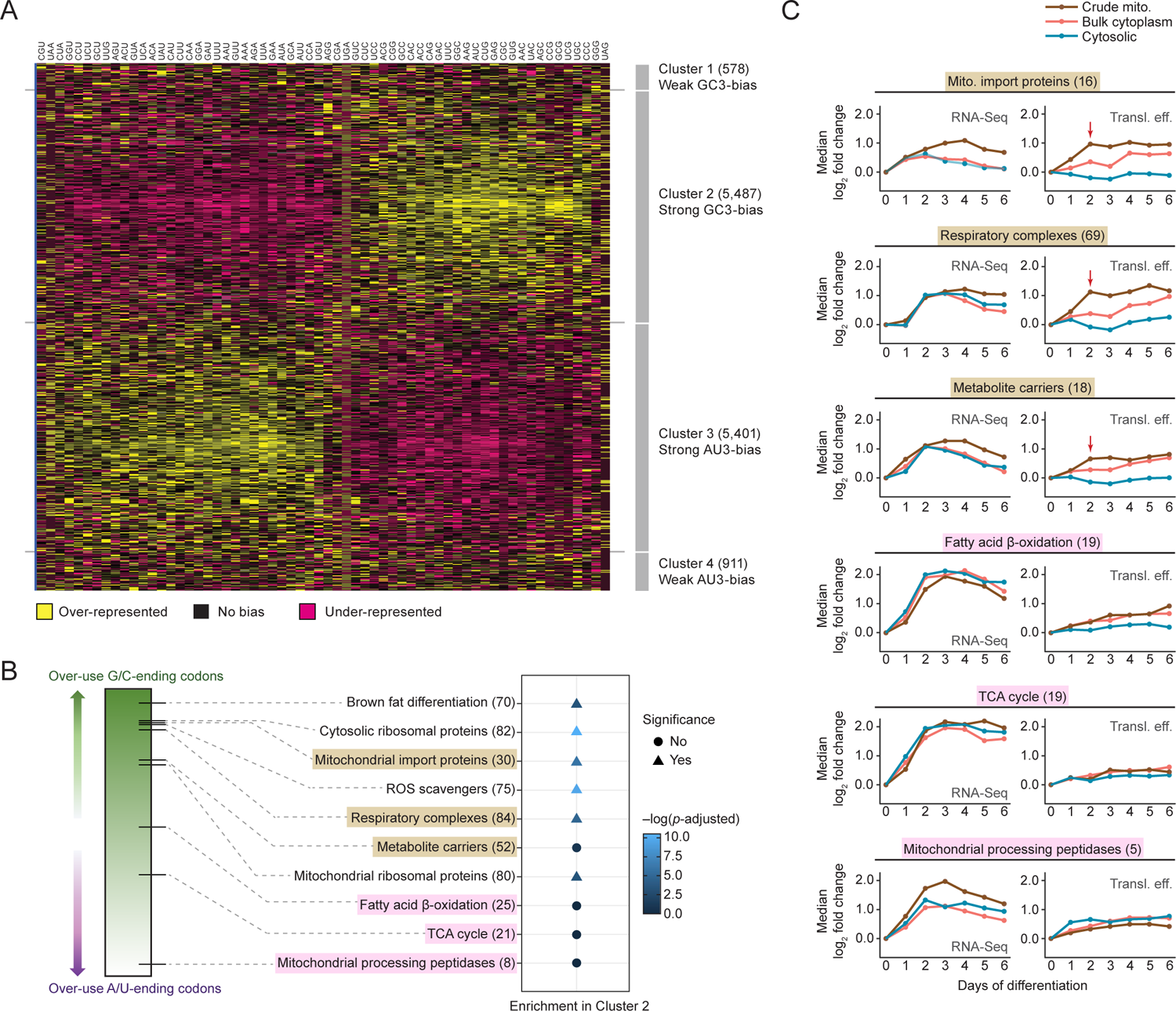
ROS-induced translation bias towards G/C-ending codons benefits mRNAs encoding brown fat differentiation and mitochondrial components, thus driving brown adipogenesis. (A) Heat map showing differential synonymous codon usage in mouse genes. Codon over- or under-representation in each ORF in the mouse genome was calculated using a Z-score matrix (see Methods). Only genes with at least one codon that has a Z-score > 2 or < –2 are included in the heat map. GC3 codon-enriched (Clusters 1 and 2) and AU3 codon-enriched (Clusters 3 and 4) groups of genes are indicated on the right. (B) Usage of GC3 codons in different functional gene groups. Genes in the mouse genome were ranked according to their aggregate Z-scores on codon usage bias. Genes in different functional groups were then placed on the hierarchy of GC3 codon usage based on the median aggregate Z-score of each group. Different functional groups were tested using the Fisher’s exact test to determine if they are significantly enriched in the strong GC3-bias cluster (Cluster 2) defined in **(A)**. Mitochondrial membrane components (brown) and mitochondrial matrix components (pink) are highlighted. The number of genes in each group is indicated in parentheses. *p* < 0.05 indicates statistical significance. (C) Expression of genes in different functional gene groups. For each functional group, the median log_2_ fold changes in RNA-Seq and translational efficiency for each day were plotted. Gene groups are highlighted as in **(B)**. Mitochondrial membrane components exhibit a boost in translational efficiency on Day 2 (red arrows) in the crude mitochondria fraction.

Although transcripts that over-use GC3 codons will benefit from the translation bias, any translational upregulation must constitute a substantial portion of the overall increase in gene expression before it can be described as a driver of brown adipogenesis. To determine if translational upregulation is a major contributing factor of expression increase for functionally relevant mRNAs, we compared changes in RNA-Seq and translational efficiency over the differentiation time course. For mRNAs encoding mitochondrial membrane components, changes in translational efficiency comprises ∼50% of the overall increase in gene expression, with the remaining half coming from increases in RNA abundance (Figure 5C). In particular, the increase in translational efficiency comes mostly from the crude mitochondria fraction, with a strong boost seen in Day 2, which is exactly when translational regulation starts to kick in in the crude mitochondria fraction once differentiation is induced (Figure 1C). Conversely, mRNAs encoding matrix components, which do not over-use GC3 codons (Figures 5B and S4A), are upregulated mainly at the RNA level (Figure 5C), with translational changes contributing ≥30% of the overall increase in gene expression. Thus, ROS-induced translation bias plays a substantial role in driving the synthesis of mitochondrial components, especially mitochondrial membrane components.

An intriguing observation is that mRNAs encoding ROS scavengers are also enriched in Cluster 2 and rank highly on the hierarchy of GC3 codon usage (Figures 5B and S4A). ROS scavengers would thus benefit strongly from the ROS-induced translation bias. This could be a means to exert negative feedback on the system to minimize the oxidative stress that will inevitably arise from the massive mitochondrial biogenesis that occurs during brown fat differentiation (see Discussion).

Besides codon usage, cellular localization also determines how mRNAs respond to the translation bias. Because we had performed RNA-Seq on different cellular compartments, we are able to identify mRNAs that are enriched in, or depleted from, the vicinity of mitochondria over the differentiation time course (Figure S5A). Analysis of mRNA enrichment ratios indicates that mRNAs that are enriched in the crude mitochondria fraction tend to have a higher percentage of GC3 codons. This is the case even before differentiation is induced (Figure 6A). The fact that these enriched mRNAs also have a higher GC3 codon content implies that they are poised to benefit from the ROS-induced translation bias, which starts off first in the mitochondrial vicinity (Figures 2C and 2D). Indeed, enriched mRNAs are translationally upregulated in the crude mitochondria fraction during differentiation (Figure 6B). In contrast, depleted mRNAs have a lower GC3 codon content and are translationally downregulated (Figures 6A and 6B).

**Figure 6.**
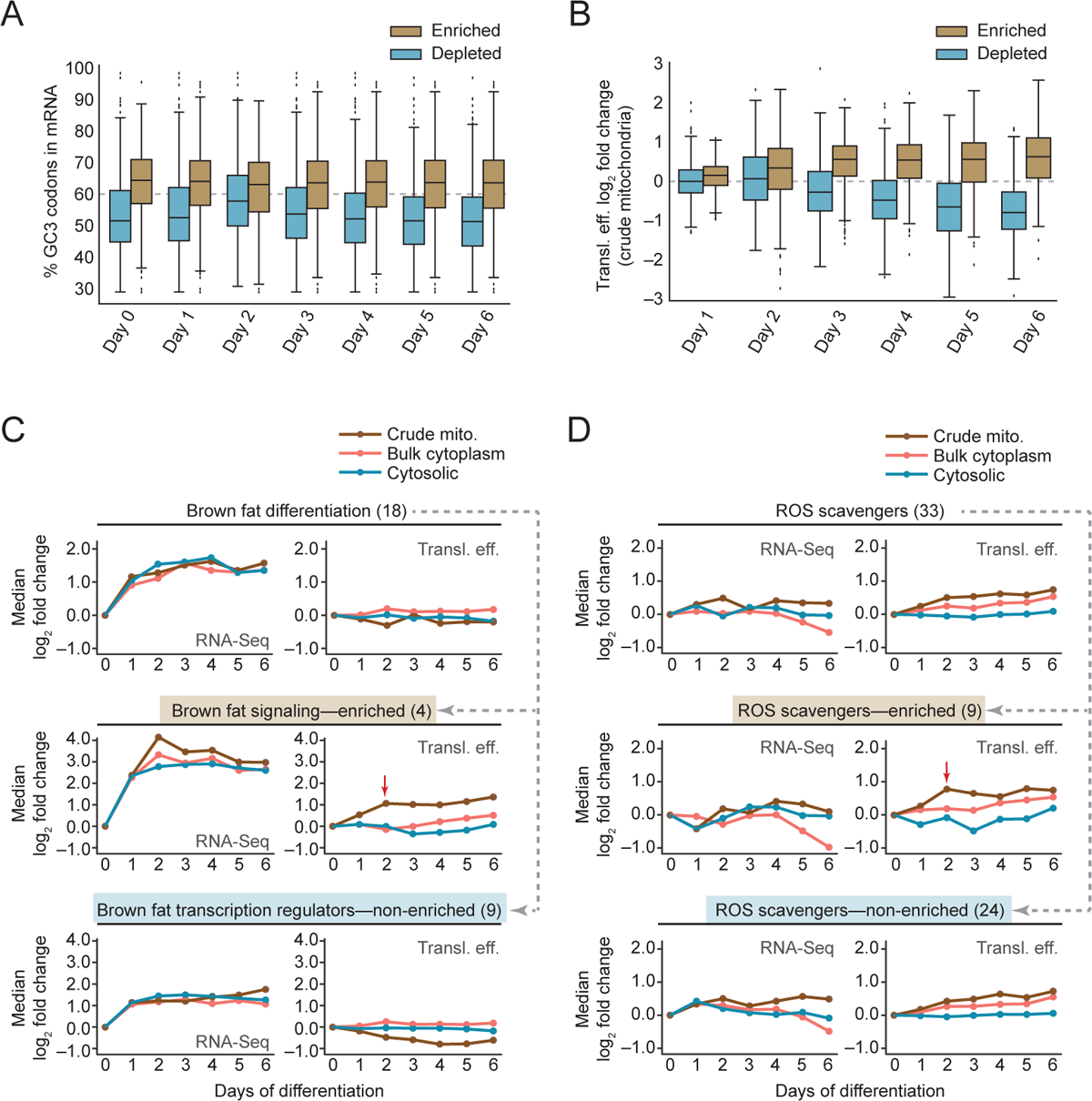
mRNAs enriched in the vicinity of mitochondria are poised to benefit from the translation bias towards G/C-ending codons early on in differentiation. (A) GC3 codon content in enriched and depleted mRNAs. The percentage of GC3 codons in each ORF was plotted, for mRNAs that were enriched in (log_2_ enrichment ratio ≥ 1), or depleted from (log_2_ enrichment ratio ≤ –1), the crude mitochondria fraction, for each day of differentiation. (B) Changes in translational efficiency in the crude mitochondria fraction for enriched and depleted mRNAs. Log_2_ fold changes in translational efficiency for each day (relative to Day 0) were plotted for mRNAs that were enriched in, or depleted from, the crude mitochondria fraction. (C) Expression of genes encoding brown fat differentiation regulators. The median log_2_ fold changes in RNA-Seq and translational efficiency for each day were plotted. Two subgroups of genes (“brown fat signaling” and “brown fat transcription regulators”) were then extracted and their expression changes similarly plotted. mRNAs encoding “brown fat signaling” components are enriched in the crude mitochondria, whereas mRNAs encoding “brown fat transcription regulators” are not enriched (see text). Only the “brown fat signaling” subgroup exhibited a boost in translational efficiency on Day 2 (red arrow) in the crude mitochondria fraction. (D) Expression of genes encoding ROS scavengers. Median log_2_ fold changes in RNA-Seq and translational efficiency for each day were plotted as in **(C)**. ROS scavenger-encoding mRNAs that were enriched in the crude mitochondria fraction (log_2_ enrichment ratio ≥ 1) were extracted as a subgroup, and their expression changes similarly plotted. The remaining mRNAs (non-enriched) were plotted as a separate subgroup. Only the enriched subgroup exhibited a boost in translational efficiency on Day 2 (red arrow) in the crude mitochondria fraction.

Previous proximity-specific expression studies have found that mRNAs encoding mitochondrial components tend to be localized to the mitochondrial surface^10–12^. Such localization is thought to facilitate rapid responses involving the mitochondria. To determine whether this is the case during brown adipogenesis, when large-scale mitochondrial biogenesis is taking place, we examined the profile of enriched/depleted mRNAs, according to the cellular localization of the proteins they encode. Inherently, crude mitochondria preparations contain endoplasmic reticulum (ER). We thus expect mRNAs encoding ER components—and components that pass through the endomembrane system—to be enriched in the crude mitochondria fraction because these mRNAs would be translated on ER-associated ribosomes. Indeed, the majority of enriched mRNAs encode ER/secretory components (Figure S5B). Other mRNAs, i.e. those that encode mitochondrial, cytoplasmic, and nuclear components, are expected to be translated on cytosolic ribosomes. We analyzed mitochondrial component-encoding mRNAs as a separate group, while all other remaining mRNAs are grouped together as “cytoplasmic/nuclear”. Consistent with previous studies, mRNAs encoding mitochondrial components tend to be enriched in, rather than depleted from, the crude mitochondria fraction. In contrast, mRNAs encoding cytoplasmic/nuclear components tend to be depleted (Figure S5B). Regardless of the eventual cellular localization of encoded proteins, enriched mRNAs—with their higher GC3 codon content— tend to be translationally upregulated in the crude mitochondria fraction, unlike depleted mRNAs (Figure S5C). These differences in GC3 codon content and mRNA localization explain why mRNAs encoding ER/secretory and mitochondrial components are translationally upregulated in the crude mitochondria fraction, whereas those encoding cytoplasmic/nuclear components are translationally downregulated (Figure S5D). Such translational upregulation would directly support ER and mitochondrial functions, which are known to play critical roles in brown adipocytes^25^. The importance of cellular localization is also reflected in two groups of mRNAs—those encoding brown fat differentiation regulators and ROS scavengers (Figure 5B)—that over-use GC3 codons, but do not encode mitochondrial components. Brown fat differentiation pathway components comprise a mix of signaling molecules, such as adipokines, and transcription factors. When we extracted the brown fat signaling components as a subgroup, it is obvious that these mRNAs—which are translated on ER-associated ribosomes and are thus enriched in the crude mitochondria fraction—exhibit the characteristic boost in translation in the crude mitochondria fraction on Day 2 (Figure 6C). In contrast, mRNAs encoding transcription factors do not exhibit this behavior. Nevertheless, ∼30% of the overall expression increase for brown fat differentiation pathway components can be attributed to translational regulation. Similarly, mRNAs encoding ROS scavengers can be separated into two subgroups based on whether they are enriched in the crude mitochondria fraction. The “enriched” subgroup clearly benefited strongly from the translation bias (boost on Day 2 in crude mitochondria fraction). In contrast, the “non-enriched” subgroup showed a more gradual increase in translational upregulation, which is consistent with the characteristics of these mRNAs— these non-enriched mRNAs do not get a translation boost in the mitochondrial vicinity on Day 2, but because they are rich in GC3 codons (Figure 5B), they eventually benefit from the translation bias after the phenomenon has spread to the rest of the cell (Figure 6D). It is worth noting that ∼60% of the upregulation in expression for ROS scavenger-encoding mRNAs can be attributed to changes in translational efficiency. ROS-induced translation bias thus contributes substantially towards upregulating the expression of ROS scavengers.

## Discussion

It was previously shown that ROS are required for white adipocyte differentiation from mesenchymal stem cells^7^. Brown adipose tissue thermogenesis is also activated by ROS through UCP1 sulfenylation^8^. We have now shown that ROS are required for brown adipogenesis. This requirement is at least in part conferred by changes in translation through ROS-induced changes in ELP3 activity. When preadipocytes are induced to differentiate, the initial mitochondrial biogenesis raises the concentration of ROS in the vicinity of mitochondria. This reduces the activity of ELP3 enzyme first in the mitochondrial vicinity, leading to a local increase in the proportion of tRNAs that are unmodified at position 34. tRNAs unmodified at position 34 do not exhibit preferential recognition of A-ending codons (Figure 3A); this in turn produces a translation bias towards mRNAs with more G-ending codons in the vicinity of mitochondria. Whether this phenomenon is mediated through all, or a subset of, ELP3-sensitive tRNAs remains to be investigated. As differentiation progresses, mitochondrial biogenesis increases and mitochondria occupy more cellular volume; this phenomenon then spreads to the rest of the cell (Figure 7A). As such, mRNAs with more GC3 codons, such as those encoding brown fat differentiation pathway components and mitochondrial components, are upregulated translationally, thus driving differentiation.

**Figure 7.**
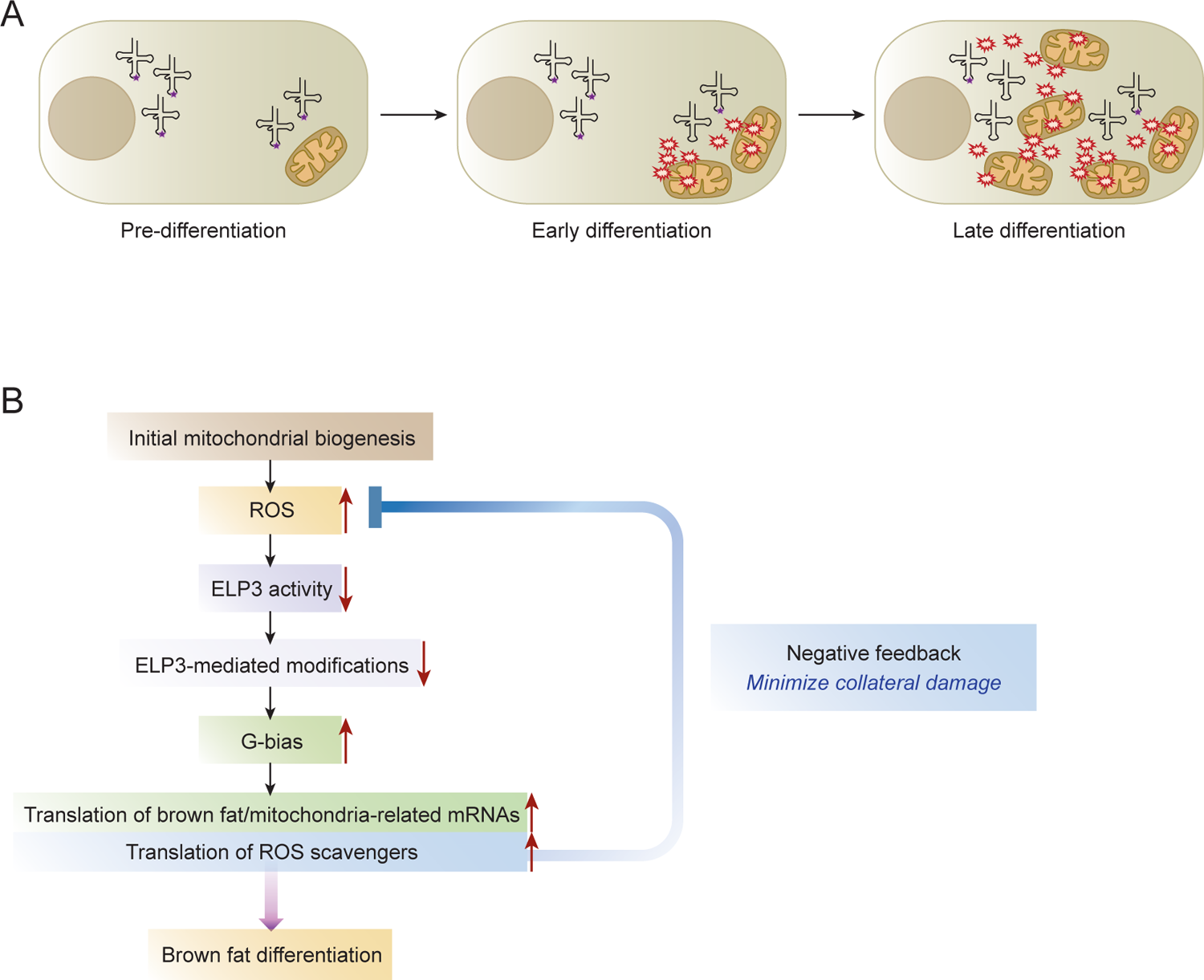
Spatiotemporal regulation of translation induced by ROS during brown adipogenesis. **(A)** Schematic showing how ROS production from mitochondrial biogenesis leads to changes in tRNA modifications through influencing the activity of ELP3. ELP3-mediated modifications are represented by purple stars. **(B)** Flow chart depicting how ROS-induced translation bias can drive brown adipogenesis and enable sustained mitochondrial biogenesis. Increased translation of ROS scavenger-encoding mRNAs can exert a negative feedback loop, thus minimizing collateral damage from oxidative stress.

However, an oxidative stress conundrum must be resolved. As massive mitochondrial biogenesis ensues during brown adipogenesis, ROS levels inevitably rise. How does the cell reconcile the need for ROS to drive differentiation and the equally pressing need to avoid cellular damage from excessive ROS? Having ROS scavengers benefit from the translation bias would provide a means to exert negative feedback. GC3 codons are over-represented in mRNAs encoding ROS scavengers, such that ∼60% of their overall expression increase can be attributed to translational upregulation. This enables cells to use ROS-induced translation bias to drive brown adipogenesis, and concurrently minimize collateral damage from having too much ROS (Figure 7B), which could be a strategy to enable sustained mitochondrial biogenesis. It would be interesting to see if this strategy is also used in other settings where substantial mitochondrial biogenesis takes place, e.g. during thermogenesis, the browning of white fat, and muscle differentiation, especially as it has already been demonstrated that ROS can induce these processes^8,26–28^. Translational control has often been described as being uniquely suited for responding to stresses, because changes in translational efficiency are rapid and reversible. Hence, such a mechanism might play an even more substantial role in the browning of white fat, which can be induced by stimulation such as exposure to cold, and is also reversible once the stimulus is removed^2^.

A key plank of this ROS-induced translational regulation strategy is that of localization. A previous ribosome profiling study done using whole cell lysates from brown adipose tissue found limited translational regulation affecting only mitochondrial components^9^. Our observations—that the bulk of translational regulation early in differentiation occurs in the mitochondrial vicinity and not in the cytosolic fraction—thus speak to the importance of analyzing different cellular compartments when studying translational regulation. It is increasingly appreciated that localized translation plays important roles in different biological contexts, enabling the cell to synthesize proteins at the locations where they are most needed and thus rapidly finetune responses. Often, such spatial regulation is orchestrated by RNA binding proteins^14^. In this study, we not only observe the enrichment of mRNAs encoding mitochondrial components in the mitochondrial vicinity, but we have uncovered a novel facet of localized translation—in addition to mRNA localization, spatiotemporal regulation of the translation bias is achieved by the localized release of a signal (ROS from mitochondria) that alters the activity of a tRNA modification enzyme (ELP3) to favor the translation of functionally relevant mRNAs that have the requisite codons (GC3 codons). Spatial partitioning of mRNAs thus synergizes with ROS-induced translation bias to ensure that the mRNAs that stand to benefit the most from the translation bias are also the ones that need to be upregulated the most. The large-scale mitochondrial biogenesis that occurs during brown fat differentiation produces mitochondria that are much more cristae-dense, compared to those seen in typical cells^4^. Hence, the need to swiftly upregulate expression of mitochondrial membrane components is much larger, compared to matrix components. It is thus intriguing to see that mRNAs encoding mitochondrial membrane components are the ones that are enriched in the mitochondrial vicinity and GC3 codon-rich. They are thus poised to benefit from the translation bias early on in differentiation. This contrasts with mRNAs encoding matrix components, which tend to under-use GC3 codons instead (Figure 5B). Another group of mRNAs that are poised to benefit from the ROS-induced translation bias are the ER-translated mRNAs. Strong mitochondria-ER contact sites have been observed in brown adipose tissue^25^. These contact sites are known to be important sites of ROS crosstalk^29^, and disrupting them in mouse preadipocytes impairs brown adipogenesis^30^. However, proximity between the ER and mitochondria necessarily means that ER components would be among the first to be exposed to, and potentially damaged by, mitochondria-produced ROS. It is thus worth noting that most of the ROS scavenger-encoding mRNAs that are enriched in the mitochondrial vicinity encode ER-resident ROS scavengers (Table S1). This would enable the cells to harness the ROS-induced translation bias to upregulate ROS scavengers at the location where they are needed most. In this way, ROS both help to fuel differentiation and mitigate potential oxidative damage.

When cells differentiate, new transcripts are needed to establish a new identity. Transcriptional changes are thus often the main driving force of differentiation. Here, we see that translational changes not only are able to drive differentiation, but they also play supportive roles. Cells that are induced to differentiate drastically upregulate the abundance of mRNAs that encode the proteins needed for the new cellular state. This necessarily means that transcripts from the previous cellular state would be present at a lower proportion in terms of relative abundance. However, certain proteins performing “housekeeping” roles are still needed to support the differentiation program. This could be why mRNAs encoding cytosolic ribosomal proteins over-use GC3 codons (Figure 5B), unlike other cytosolic component-encoding mRNAs (Figure S5D). This feature could ensure that the translation machinery is maintained at a capacity that is high enough to synthesize the new proteins needed for differentiation. A second case in point are the ROS scavengers. ROS scavenger-encoding mRNAs either maintain, or are reduced in, their relative abundance during differentiation (Figure 6D). However, because these mRNAs over-use GC3 codons, they are translationally upregulated, and are thus able to perform the critical role of minimizing oxidative damage during brown adipogenesis.

Translational reprogramming due to codon usage biases and tRNA modifications have previously been reported to enable cells to adapt to stresses. In yeast, tRNA methyltransferases Trm9 and Trm4 catalyze different modifications on position 34, and in so doing, alter wobble base recognition, leading to selective translation of mRNAs that encode stress-responsive proteins^19,31^. Here, we report that ELP3 is sensitive to ROS changes by virtue of its Fe-S cluster, and that this effect is large enough to cause a translation bias towards G-ending codons and help drive a differentiation program. This sensitivity to ROS is reminiscent of the rRNA and tRNA modification enzyme, RlmN, in *Enterococcus faecalis*. RlmN activity is downregulated through ROS-mediated inactivation of the enzyme’s Fe-S cluster, leading to codon-biased proteomic changes that enable the bacteria to mount a stress response program^32^. Understanding whether other Fe-S cluster-containing modification enzymes^33^ behave similarly during brown adipogenesis would be an important next step. Interestingly, ELP3 has been found to localize to mitochondria in HeLa cells^34^ and *Toxoplasma gondii*^35^. It was also recently reported that a novel splice variant of ELP3 is localized to the mitochondrial matrix, where it modifies mitochondrial tRNAs, and in so doing, protects the tRNAs from degradation^36^. If these mechanisms are also in play during brown adipogenesis, they will add a further dimension to how translational changes might be localized and effected.

The fact that codons can influence translation implies that the elongation step must be affected. Oxidative stress can slow down translation elongation through various means^37^. However, in this study, initiation is still the rate-limiting step because the translation bias we detected from sequencing was reproduced in the luciferase reporter assays, i.e. having more GC3 codons genuinely led to more protein synthesis—an effect that cannot be produced by a slower elongation rate. Hence, just as codon optimality has been reported to impact mRNA stability through as yet incompletely defined mechanisms^38^, codon optimality impacts the translation initiation rate of mRNAs, through an as yet undefined link to translation elongation.

It was previously shown that the composition of tRNA pools in proliferating cells differs from that present in differentiating cells, and that these tRNA changes are coordinated with changes in mRNA expression^39^. Here, we report that the composition of tRNAs changes, not because of abundance changes, but because of ROS-induced changes in tRNA modification activity. This, coupled with the spatial partitioning of mRNAs, enables changes in the tRNA pool to be harnessed at the right time and at the right location to drive brown adipogenesis. A requirement for this model (Figure 7A) is that the proportion of unmodified tRNAs must increase as differentiation progresses; in other words, there must be timely turnover of the pre-existing modified tRNAs. tRNAs are known to have long half-lives, ranging from tens of hours to days^40^. However, intriguingly, tRNA stability can be modulated by modifications and can also be reduced under oxidative conditions. In particular, thiolated tRNAs—which include ELP3-modified tRNA-Lys(UUU), tRNA-Gln(UUG), tRNA-Glu(UUC)—can be dethiolated under oxidative conditions, leading to cleavage and/or altered base-pairing^41^. Whether the ROS produced during brown adipogenesis exert additional impact through modulating the stability of modified tRNAs would be an important question to address.

While excessive ROS is highly damaging to cells, it is increasingly appreciated that ROS at physiological concentrations play critical cellular roles, including as signaling molecules^42^. In other words, ROS have a hormetic effect on cells—both too much or too little can be detrimental. Here, we show that ROS-induced translation bias towards GC3 codons play a critical role in driving brown adipogenesis, enabling massive mitochondrial biogenesis while minimizing oxidative damage. These data lay out, at least in part, the molecular basis for ROS hormesis, and lend insights towards understanding the utility of antioxidants as therapeutics. Antioxidants have been proposed to treat myriad disorders, but many clinical trials employing antioxidant supplementation have thus far not shown much benefit; some have even reported detrimental effects^43,44^. The molecular consequences described in this study could explain how blanket usage of antioxidants can negate the beneficial effects of ROS. Analogously, mitohormesis—a phenomenon in which mild mitochondrial stress can lead to beneficial effects^45^—might employ similar mechanisms. It was previously reported that transient oxidative stress can be induced in a mouse model that enables reversible knockdown of the mitochondrial superoxide dismutase SOD2, leading to an adaptive, sustained response in mitochondrial biogenesis and antioxidant upregulation in a cell type-specific manner, which cannot be fully explained by known transcriptional networks^46^. Our findings, that different mRNAs can be translated differently in response to ROS, could explain these molecular changes, and why responses might differ in different tissue/cell types, which inherently have different levels of ROS exposure. This provides an additional framework to understand these molecular effects and could shed new light on the mechanisms that mediate mitohormesis. Lastly, GC3 codon usage bias is conserved between mouse and human^47^. Coupled with the observation that cancerous and normal tissues use GC3 codons differently^39,48^, the phenomenon we observed might apply beyond metabolic disorders. Therefore, our findings could have important implications beyond contexts that involve obvious mitochondrial changes.

## Methods

### Brown adipocyte differentiation

Immortalized brown preadipocytes from mouse origin^15^ were a gift from Dr. Lei Sun. For differentiation into mature brown adipocytes, cells were cultured in high-glucose DMEM supplemented with 870 nM insulin (Sigma) and 2 nM triiodothyronine (Sigma) [differentiation medium] for three days to reach full confluency. Differentiation was then induced (Day 0) with the additional supplementation of an induction cocktail [250 μM isobutylmethylxanthine (Sigma), 0.5 μM dexamethasone (Sigma) and 60 μM indomethacin (Sigma)]. Two days after induction, cells were switched back to differentiation medium (Day 2). Medium was changed every two days and full differentiation was achieved on Day 6. All experiments were performed with cells at passages 3 to 5.

### Cell harvesting and cellular fractionation

Cells were treated with cycloheximide (100 μg/mL, 10 min) and chloramphenicol (100 μg/mL, 20 min) to arrest cytosolic and mitochondrial translation, respectively. Cycloheximide (100 μg/mL) and chloramphenicol (100 μg/mL) were subsequently included in all buffers used for harvesting and fractionation. After translation arrest, cells were trypsinized and pelleted at 300 x g, for 5 min. For each day of differentiation, cells from multiple 15-cm dishes were combined and then split into portions for harvesting the bulk cytoplasm, crude mitochondria, or the cytosolic fraction without mitochondria. To obtain the bulk cytoplasm, pelleted cells were lyzed in ice-cold lysis buffer [10 mM Tris-HCl, pH 7.4, 5 mM MgCl_2_, 100 mM KCl, 1% Triton X-100, 2 mM dithiothreitol, 100 U/mL RNasin (Promega), complete protease inhibitor (Roche)]. The lysate was homogenized 10 times with a 26-gauge needle at 4 °C and centrifuged at 1,300 x g for 10 min to remove the nuclear pellet. The resulting supernatant was saved as the bulk cytoplasm. To obtain the crude mitochondria and cytosolic fractions, pelleted cells were resuspended in ice-cold RSB hypotonic buffer (10 mM Tris-HCl, pH 7.5, 10 mM NaCl, 1.5 mM MgCl_2,_ complete protease inhibitor), and allowed to swell for 5 min on ice. The cell suspension was then homogenized with a 7-mL glass homogenizer. Efficiency of cell lysis was checked by light microscope. The mannitol and sucrose concentrations of the buffer were then adjusted to 210 mM and 70 mM respectively by adding 2.5 x MS homogenization buffer (12.5 mM Tris-HCl, pH 7.5, 525 mM mannitol and 175 mM sucrose, complete protease inhibitor). Unlyzed cells and nuclei were then pelleted at 1,300 x g for 3 min at 4 °C. This clarification spin step was repeated another two times. Mitochondria were then pelleted by centrifugation at 12,000 x g for 15 min at 4 °C. The resulting supernatant was saved as the cytosolic fraction. The mitochondrial pellet was washed in 1 x MS homogenization buffer (5 mM Tris-HCl, pH 7.5, 210 mM mannitol, 70 mM sucrose, complete protease inhibitor) and centrifuged at 12,000 x g for 15 min at 4 °C. The resulting pellet was then resuspended in mitochondrial lysis buffer (10 mM Tris-HCl, pH 7.5, 20 mM MgCl_2_, 100 mM KCl, 1% Triton X-100, 5 mM β-mercaptoethanol, complete protease inhibitor and 100 U/mL RNasin). The lysate was then homogenized 10 times with a 26-gauge needle and incubated on a rotator at 4 °C for 20 min to facilitate complete lysis. The lysate was then clarified by centrifugation at 12,000 x g for 15 min at 4 °C. The resulting supernatant is the crude mitochondria fraction. The bulk cytoplasm, cytosolic fraction (without mitochondria), and crude mitochondria fraction were then snap-frozen before downstream analysis.

### Oil Red O staining

Cells were washed twice in PBS and fixed with 10% paraformaldehyde. Oil Red O stock solution (0.5% w/v) was prepared by dissolving 0.5 g Oil Red O (Sigma-Aldrich) in 100 mL isopropanol. The working solution (0.3% w/v) was prepared freshly before use by diluting the stock solution with deionized water and filtering through Whatman #1 filter paper. Cells were stained with Oil Red O working solution for 1 h at room temperature and then washed at least three times with deionized water.

### MitoTracker staining

Cells were washed with PBS and incubated with pre-warmed staining solution containing 100 nM MitoTracker Red CMXRos (Invitrogen) at 37 °C for 30 min. Cells were then washed with media and fixed with 3.7% formaldehyde at 37 °C for 15 min, after which cells were washed twice with PBS, and stained with DAPI staining solution (Invitrogen) for 5 min at room temperature. Cells were then washed twice with PBS and visualized using the ZOE Fluorescent Cell Imager (Bio-Rad).

### Transfections and dual luciferase reporter assays

Brown preadipocytes were seeded in 6-well plates and induced to differentiate. Cells were transfected with plasmids carrying *Renilla* luciferase and the appropriate firefly luciferase variant (see “Firefly luciferase reporter constructs”) using jetPrime transfection reagent (Polyplus) according to manufacturer’s instructions. Transfection medium was replaced 4 h after transfection, with the appropriate culture medium (depending on the day of the differentiation protocol). Cells were harvested to assess luciferase activity 24 h after transfection. Dual luciferase reporter assays were performed using the Dual-Luciferase Reporter Assay System (Promega) according to manufacturer’s instructions. Luciferase activity was detected using the GloMax Discover Microplate Reader (Promega). All experiments were done in triplicates.

### Firefly luciferase reporter constructs

Modified versions of firefly luciferase were designed according to the following rationale. To obtain a G-maximized luciferase reporter construct, all codons that could be recognized differently due to ELP3-mediated modification had their third base modified to “G” if they did not end in a “G” originally. Conversely, to obtain an A-maximized luciferase reporter construct, all codons that could be recognized differently due to ELP3-mediated modification had their third base modified to “A” if they did not end in an “A” originally. These mutated versions of firefly luciferase were chemically synthesized (Gene Universal) and cloned into pcDNA3.1/Zeo(+) vector (Invitrogen). Successfully cloned plasmids were verified by Sanger sequencing. All three firefly luciferase reporter constructs (wild-type, G-maximized, A-maximized) have identical 5′UTR and 3′UTR sequences.

### cDNA synthesis and quantitative reverse transcription PCR (qRT-PCR)

Total RNA was treated with DNase I and purified using the RNA Clean & Concentrator kit (Zymo Research) according to manufacturer’s instructions. Purified RNA was reversed transcribed using the SuperScript VILO cDNA Synthesis Kit (Invitrogen). Quantitative PCR was performed using the PowerUp SYBR Green Master Mix (Applied Biosystems), together with the StepOnePlus Real-Time PCR System (Applied Biosystems), according to manufacturer’s instructions. All samples were run in triplicates. Primer sequences used are listed below.

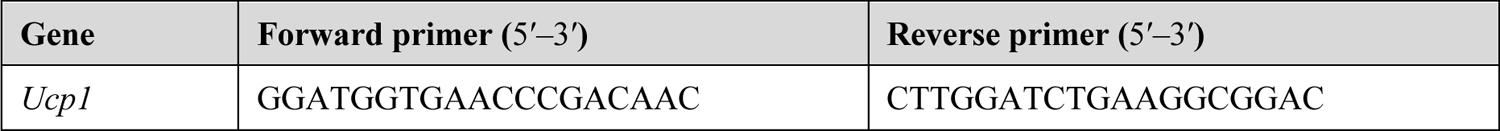
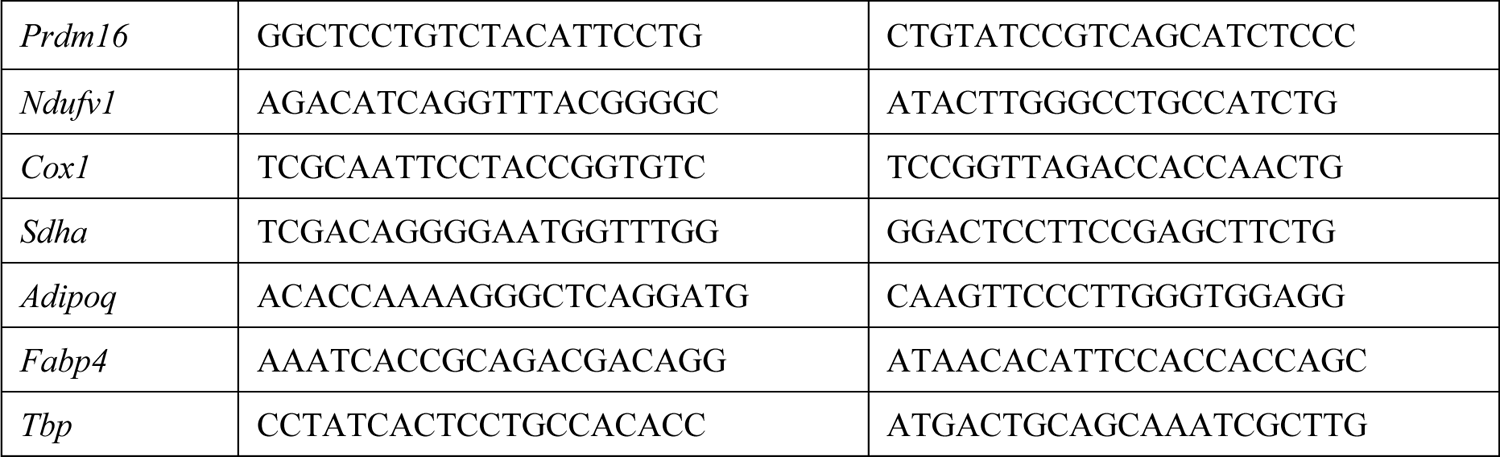

### tRNA abundance analyses

Purified total RNA samples from the crude mitochondria fraction or the cytosolic fraction from Days 0, 2, 4 and 6, were submitted to ArrayStar for relative quantification of different tRNA species by quantitative real-time PCR. Relative fold changes in tRNA abundance were derived by normalizing the measurement of a particular tRNA from each day of differentiation to the analogous measurement from Day 0. tRNAs shown in Figure S2 are named according to the nomenclature adopted by ArrayStar: https://www.arraystar.com/assets/1/6/Mouse_tRNA_full_list.pdf.

### Isolation and purification of small RNAs for tRNA modification mass spectrometry

Crude mitochondria and cytosolic fractions from Days 0, 2, 4 and 6 were obtained as described above. Total RNA was isolated using TRIzol LS reagent (Invitrogen) according to the manufacturer’s instructions. Small RNAs (17– 200 nucleotides) were extracted using the RNA Clean & Concentrator-100 kit (Zymo Research) according to manufacturer’s instructions, with an additional ethanol precipitation step to concentrate samples.

### Analysis of ribonucleosides by chromatography-coupled tandem mass spectrometry

Analysis of the ribonucleosides was performed as previously described^32^ with some modifications. Briefly, purified small RNA (5 µg) was enzymatically hydrolyzed for 6 h at 37 °C in a 50 µL digestion cocktail containing 1 U benzonase, 4 U calf intestinal alkaline phosphatase, 0.1 U phosphodiesterase I, 0.1 mM deferoxamine, 0.1 mM butylated hydroxytoluene, 5 ng coformycin, 100 nM internal standard [^15^N]_5_-deoxyadenosine, 2.5 mM MgCl_2_ and 5 mM Tris buffer pH 8.0. Samples were cleaned up using a 10 kDa cutoff filter (Nanosep). Analysis of the tRNA modifications and the four canonical ribonucleosides was conducted by liquid chromatography-coupled triple quadrupole mass spectrometry (LC-MS/MS) and liquid chromatography-coupled diode array detector (LC-DAD), respectively. The ribonucleoside mixtures were resolved on a Hypersil GOLD aQ C18 column (2.1 x 100 mm, 1.9 μm particle size) mounted on an Agilent 1290 HPLC system and linked to an Agilent 1290 Infinity DAD and an Agilent 6495 triple quadruple mass spectrometer. The column was kept at 35 °C and the auto-sampler was cooled at 4 °C. The UV wavelength of the DAD was set at 260 nm and the electrospray ionization of the mass spectrometer was performed in positive ion mode with the following source parameters: drying gas temperature 120 °C with a flow of 11 L/min, nebulizer gas pressure 20 psi, sheath gas temperature 400 °C with a flow of 12 L/min, capillary voltage 1,500 V and nozzle voltage 0 V. Compounds were quantified in multiple reaction monitoring (MRM) mode and instrument parameters were optimized using synthetic standards. Data acquisition and processing were performed using MassHunter software. The signal intensity of the modifications was normalized against the combined intensity of the four canonical ribonucleosides to correct for variation in RNA quantities.

### Measurement of reactive oxygen species

Cells were seeded in 96-well black wall/clear bottom plates. Cellular ROS and superoxide levels were assessed using the ROS/Superoxide Detection Assay Kit (Abcam) according to manufacturer’s instructions. Mitochondrial superoxide was assessed using the Mitochondrial Superoxide Assay Kit (Abcam) according to manufacturer’s instructions. Measurements were done using a fluorescence microplate reader. For each condition, six technical replicates were performed. Uninduced cells were kept in differentiation media without the induction cocktail.

### Library preparation for RNA-Seq and ribosome profiling

For each sample type (bulk cytoplasm, crude mitochondria, or cytosolic fraction), portions were set aside for RNA-Seq and ribosome profiling. For the RNA-Seq portion, RNA was extracted using TRIzol LS Reagent (Invitrogen) according to manufacturer’s instructions and purified using the RNA Clean & Concentrator kit (Zymo Research). Purified RNA was subjected to ribosomal RNA depletion using the Ribo-Zero Magnetic Gold Kit (Human/Mouse/Rat; Epicentre), according to manufacturer’s instructions. The resulting mRNA was then fragmented by partial hydrolysis in alkaline fragmentation buffer (10 mM Na_2_CO_3_, 90 mM NaHCO_3_, 2 mM EDTA, pH ≈ 9.3) at 95 °C for 20 min. For the ribosome profiling portion, RNase I (Ambion) was added to the extract, and the reaction incubated for 30 min on a shaker at 4 °C to digest polysomes. Digested lysates were loaded onto pre-cooled 10–50% sucrose gradients and centrifuged in an SW-41Ti rotor at 36,000 rpm using an Optima L-100 XP Ultracentrifuge (Beckman Coulter) at 4 °C for 2 h. Gradients were fractionated using the Gradient Station fractionator (Biocomp Instruments), with absorbance at 254 nm monitored using a Triax Flow Cell (Biocomp Instruments). Monosome fractions were pooled and treated with 200 μg/mL Proteinase K (Roche) and 1% sodium dodecyl sulfate at 50 °C for 30 min. Ribosome-protected fragments (RPFs) were then purified with acid phenol:choloform (pH 4.5, Ambion), according to manufacturer’s instructions. Fragmented RNAs (for RNA-Seq) and purified RPFs (for ribosome profiling) were then separated on a 10% polyacrylamide-urea gel. Fragments corresponding to 33–48 nucleotides (RNA-Seq) and 27‒33 nucleotides (ribosome profiling) were size-selected and purified. Purified fragments were dephosphorylated at their 3′ ends with T4 polynucleotide kinase (New England Biolabs) at 37 °C for 1 h and ligated to a 3′ adaptor sequence using T4 RNA ligase 2 (truncated K227Q, New England Biolabs) at 22 °C for 3 h. For 3′-ligated RPF fragments, an additional subtractive hybridization step was used to remove top ribosomal RNA contaminants, using biotinylated oligonucleotides that are antisense to the targeted rRNA sequences, and Dynabeads MyOne Streptavidin C1 beads (Invitrogen). Thereafter, 3′-ligated RNA fragments and subtracted 3′-ligated RPF fragments were reverse-transcribed using SuperScript III Reverse Transcriptase (Invitrogen) at 48 °C for 30 min. Barcodes for multiplexing were incorporated during reverse transcription. Template RNA was degraded by adding 100 mM NaOH and incubating at 98 °C for 20 min. The reverse-transcribed products were then gel-purified and circularized using CircLigase ssDNA Ligase (Epicentre), according to manufacturer’s instructions. Circularized products were used as templates for PCR amplification, which was carried out with Phusion High-Fidelity DNA Polymerase (Thermo Scientific). PCR products were assessed for purity and concentration on the Bioanalyzer using the High Sensitivity DNA Kit (Agilent Technologies). Samples were multiplexed and sequenced on the Illumina HiSeq 2500 system (Genome Technology Core, Whitehead Institute). Two biological replicates of each time-point (Days 0, 1, 2, 3, 4, 5 and 6) and sample type (bulk cytoplasm, crude mitochondria, or cytosolic fraction) were prepared. Antisense biotinylated oligonucleotides used to subtract the most frequent rRNA contaminants from ribosome profiling libraries are listed below.

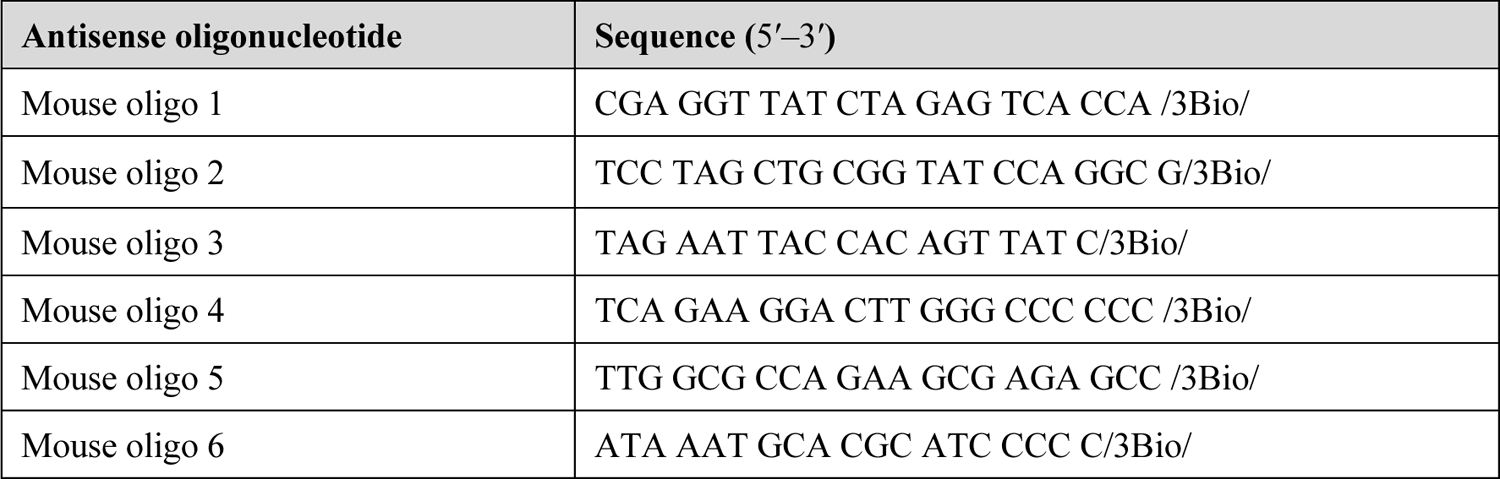
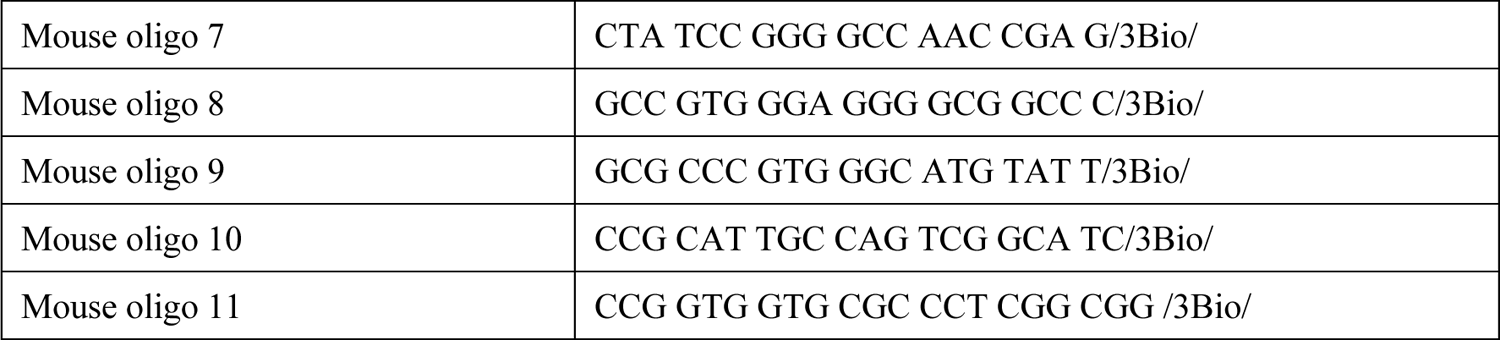

### Sequence analyses

Prior to alignment, 3’ adaptors were trimmed from the raw sequences using Cutadapt^49^. Sequencing reads were then aligned to the mouse genome (mm9) with the Bowtie short-read mapping program^50^, using the first 25 nucleotides as the ‘seed’ region. A three-stage iterative mapping was performed—reads that failed to map to the genome when no seed mismatches were allowed were fed into Bowtie again, allowing for one seed mismatch, or two seed mismatches, with each subsequent round of mapping. Reads with multiple equivalent hits to the genome were discarded, as were reads that mapped to ribosomal RNAs and transfer RNAs. To capture reads uniquely spanning splice junctions, reads that failed to map to the genome were mapped to a set of reference transcripts built based on RefSeq mRNA sequences (with the refFlat file generated on 16 June 2017, downloaded from the UCSC Genome Browser). The same three-stage iterative mapping was used and again reads with multiple equivalent hits were discarded. Reads with unique alignments from mapping to the genome or mapping to the reference transcript database were combined for subsequent analyses.

### Gene expression analyses

Differential gene expression analyses were conducted using the DESeq2 package^51^. Only genes with read counts ≥ 10 in all sample types (bulk cytoplasm, crude mitochondria, or cytosolic fraction) across all time-points were used as input. Expression fold changes of genes were defined as the expression value of any given day normalized to that of Day 0. The translational efficiency of each mRNA was derived by normalizing the number of ribosome-protected fragments to the number of RNA-Seq tags that map to the open reading frame. RNA enrichment ratio was defined as the RNA expression value in the crude mitochondria fraction normalized to that in the cytosolic fraction. An mRNA with RNA log_2_ enrichment ratio ≥ 1 is considered “enriched” in the mitochondrial vicinity; conversely, an mRNA with RNA log_2_ enrichment ratio ≤ –1 is considered “depleted”.

### Codon translation coefficient plots

The Pearson correlation coefficient between the translational efficiency log_2_ fold changes of mRNAs and the frequency of occurrence of individual codons in the open reading frames of the same mRNAs was first derived for every coding codon. This correlation coefficient was defined as the codon translation coefficient (CTC). The CTCs for all 61 coding codons were then plotted in a consolidated bar plot for each time point.

### Gene-specific codon usage analyses

Gene-specific codon usage patterns were determined as previously described^23,24^. Briefly, the gene-specific codon frequency values were determined for all genes in the mouse genome (mm9). For each codon, a genome-average codon frequency value was also derived from all the 20,485 genes in the genome. A gene-specific codon Z-score was then calculated as such: *Z* = (*G* – *A*)/*S*, where *G* is the actual codon frequency for a particular gene, *A* is the genome-average codon frequency for the particular codon in question, and *S* is the corresponding standard deviation for each codon from all genes in the reference genome. Hierarchical clustering and heat map analyses were performed using Cluster and TreeView^52–54^. Only genes with at least one codon Z-score > 2 or < –2 were included in the heat map in Figure 5A (12,377 genes). To rank genes according to their GC3 codon bias, codon-specific Z-scores were summed for each gene. As such, the heat map contains 12,377 genes but the one-dimensional aggregated Z-score ranking hierarchy in Figure 5B includes all genes in the reference transcript database (20,485 genes) used in this study. The Fisher’s exact test was used to determine statistical enrichment of different functional gene groups in specific codon usage clusters. Clusters were demarcated based on the heat map in Figure 5A.

### Annotations

A non-redundant list of mitochondrial proteins was curated using the MitoCarta2.0^55^ and COMPARTMENTS^56^ databases, with cross-checking from the literature. mRNAs encoding ER-resident proteins, and proteins that would pass through the endomembrane system, are expected to be translated on ER-associated ribosomes, and are thus expected to be enriched in the crude mitochondria fraction, which contains mitochondria and ER. A list of “ER/secretory” proteins were thus curated using SignalP 4.0^57^ (proteins predicted by SignalP to have the signal peptide would be included) and the COMPARTMENTS database. All other genes (not included in the “Mitochondrial” or “ER/secretory” categories described above) were classified as “Cytoplasmic/nuclear”. Other functional gene groups used in this study were curated using the KEGG database^58^, with cross-checking from the literature.

## Acknowledgements

We thank P. Matsudaira, Y.X. He, L. Sun, and S. Pervaiz for advice and discussions, and the Whitehead Institute’s Genome Technology Core for sequencing. The immortalized mouse brown preadipocyte cell line was a gift from Dr L. Sun. This work was supported by the Institute of Molecular and Cell Biology, Agency for Science, Technology and Research (A*STAR), Singapore, research grants from the Biomedical Research Council, A*STAR, Singapore [grant number BMRC/YIG/13/1/01YA/001 (to H.G.)], the National Medical Research Council, Singapore [grant numbers NMRC/OFYIRG/0019/2016, NMRC/OFIRG/0015/2016, and NMRC/OFIRG/0038/2017 (to H.G.)], the National Research Foundation of Singapore through the Singapore-MIT Alliance for Research and Technology Antimicrobial Resistance Interdisciplinary Research Group (to P.C.D.), and the USA National Institutes of Health [grant numbers ES026856, ES031529, and ES024615 (to T.J.B.)]. J.Y.I. was supported by A*STAR and the NUS Graduate School for Integrative Sciences and Engineering. Y.L. was supported by A*STAR.

## Author Contributions

J.Y.I., I.W., P.C.D., and H.G. contributed to the design of the study. J.Y.I., I.W., L.T.L., Y.L., D.E.-J.T., C.S.C.C., L.C., and T.J.B. performed the experiments and analyzed the data, with input from the other authors. All authors contributed to the preparation of the manuscript.

## Author Information

Sequencing data were deposited in the Gene Expression Omnibus (www.ncbi.nlm.nih.gov/geo/) under accession number GSE249222.

## Supplemental Figure legends

**Figure S1.**
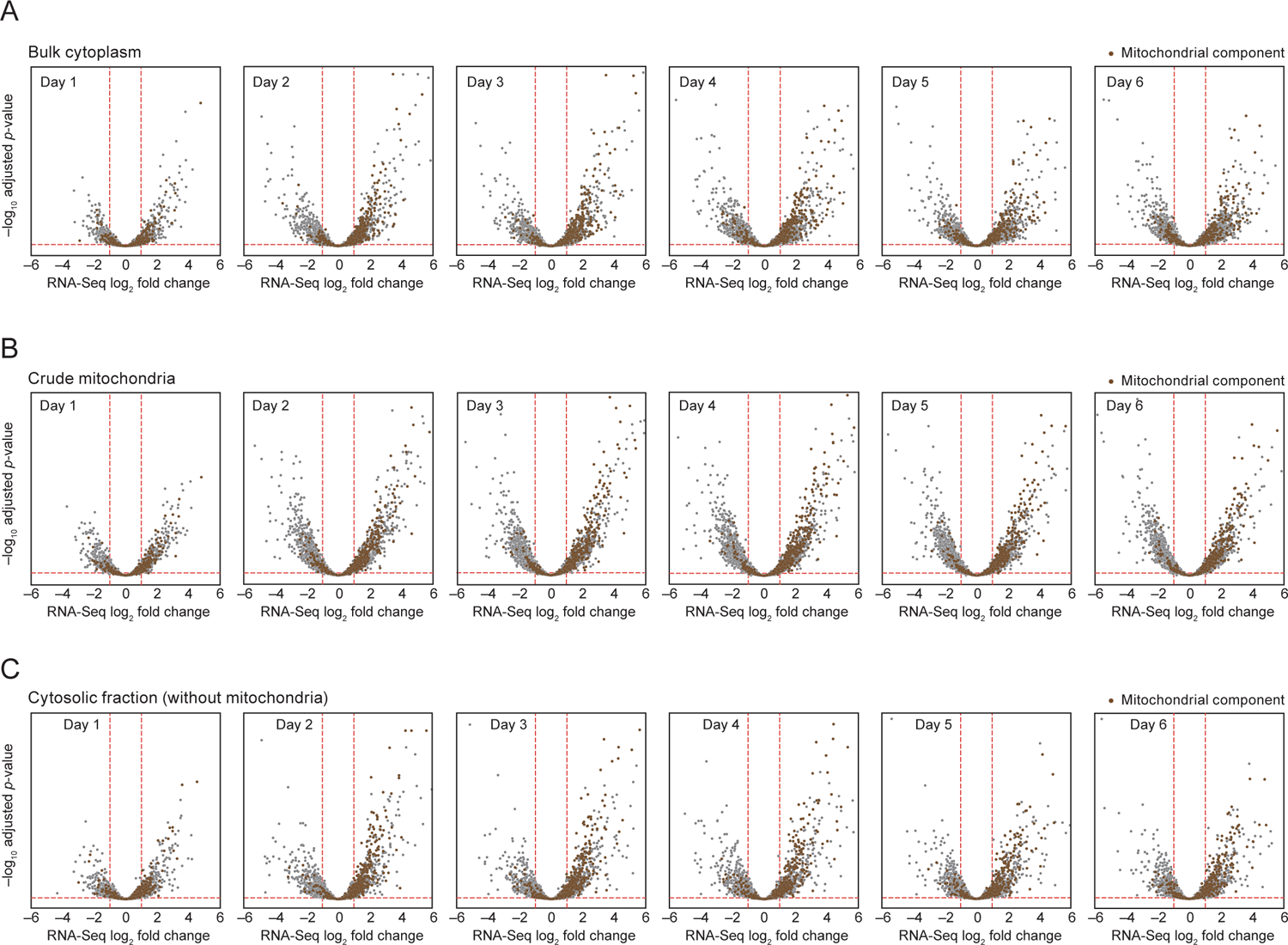
RNA expression changes take place in different cellular compartments once brown adipogenesis is induced, related to **Figure 1**. (A) Volcano plots showing RNA-Seq fold changes in the bulk cytoplasm for each day (relative to Day 0) as brown adipogenesis progresses. Genes encoding mitochondrial components are highlighted in brown. (B) RNA-Seq fold changes in the crude mitochondria fraction. Otherwise, as in (A). (C) RNA-Seq fold changes in the cytosolic fraction. Otherwise, as in (A).

**Figure S2.**
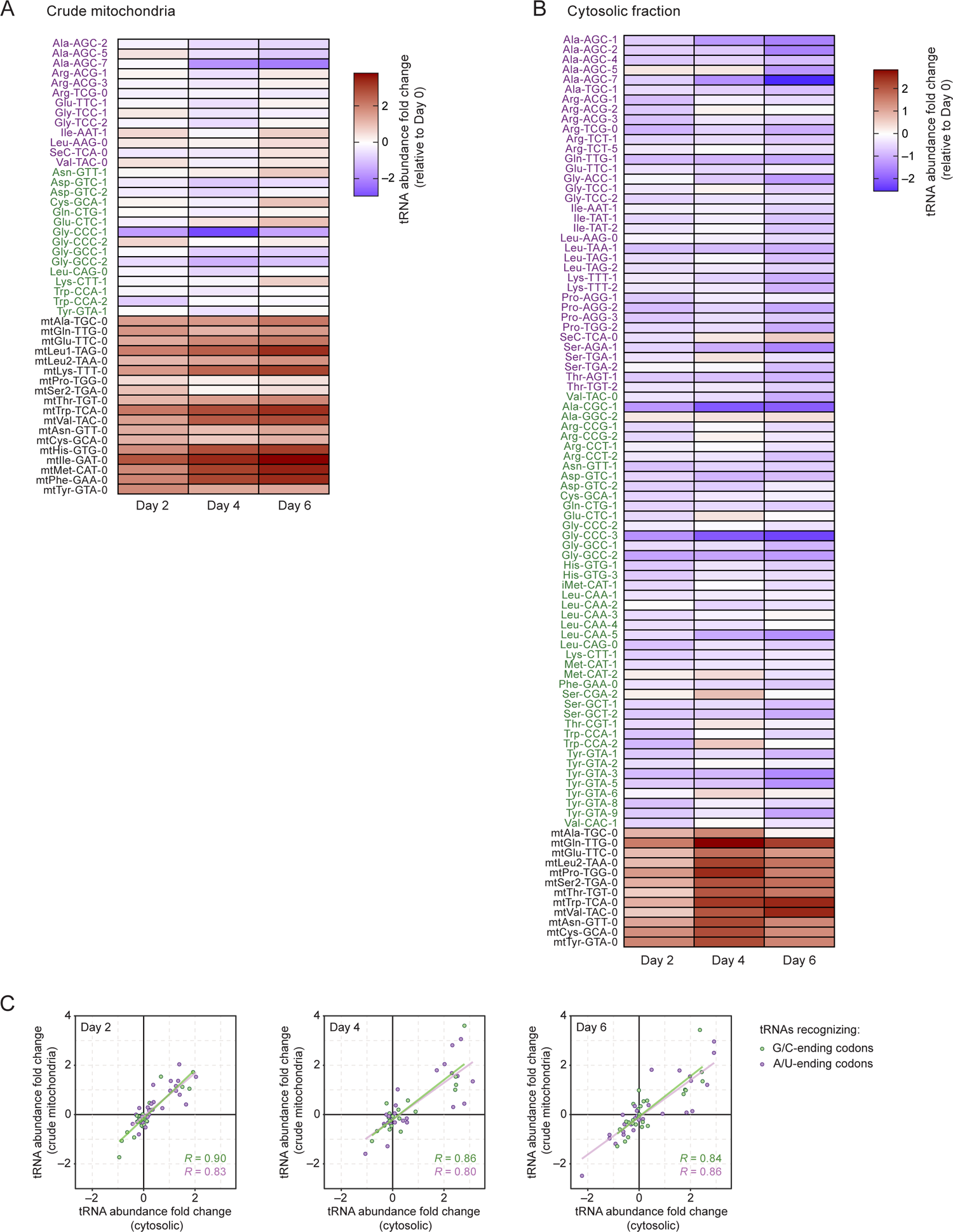
Changes in tRNA abundance do not explain the observed translation bias towards G/C-ending codons, related to Figure 3. **(A)** Heat map showing changes in tRNA abundance in the crude mitochondria fraction during brown adipogenesis. tRNA abundance levels were measured by qRT-PCR and plotted relative to abundance levels in Day 0. Cytosolic tRNAs are ordered according to whether their cognate codons are GC3 codons (green) or AU3 codons (purple). Mitochondrial tRNAs (black) are included. Data shown is the average of two biological replicates. (B) Heat map showing changes in tRNA abundance in the cytosolic fraction. Otherwise, as in (A). **(C)** Correspondence between fold changes in tRNA abundance in the crude mitochondria and cytosolic fractions. tRNAs that recognize GC3 codons (green) or AU3 codons (purple) are highlighted. The Pearson correlation coefficients for all groups of tRNAs are indicated.

**Figure S3.**
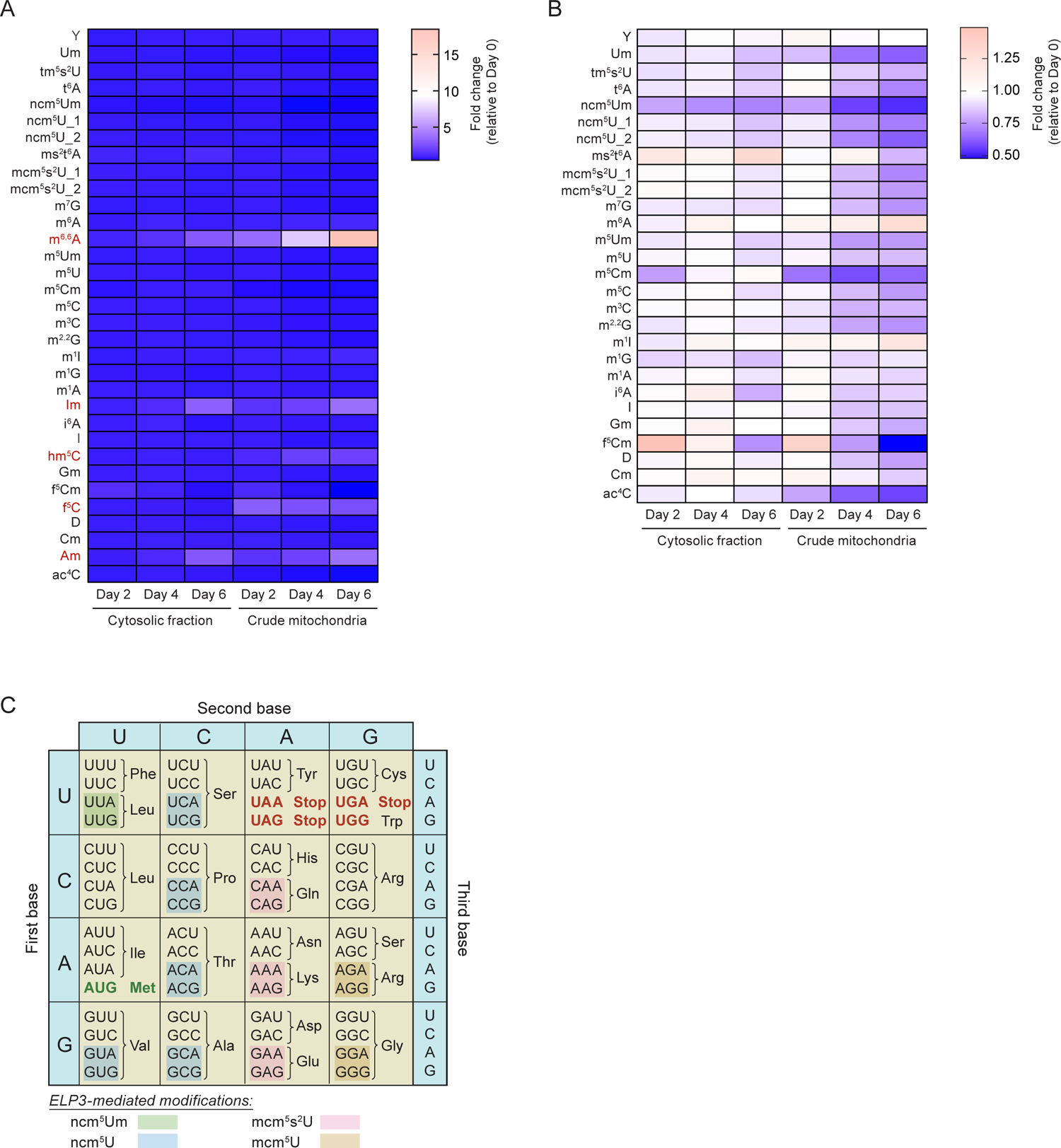
Changes in tRNA modifications, due to reduced ELP3 activity, could explain observed translation bias towards G/C-ending codons, related to Figure 3. (A) Heat map showing changes in tRNA modifications (relative to Day 0) during brown adipogenesis. tRNAs were isolated from the crude mitochondria and cytosolic fractions from Days 0, 2, 4 and 6 of differentiation, and tRNA modifications were assessed using mass spectrometry. Where indicated, “_1” and “_2” denote two unique transitions on the mass spectrometer. Five modifications exhibited outsized fold changes (highlighted in red), masking the changes of other tRNA modifications. (B) Heat map showing changes in tRNA modifications (relative to Day 0) during brown adipogenesis, without the five outlier modifications described in (A). (C) Genetic code table, with the codons affected by ELP3-mediated modifications highlighted. ELP3, as part of the Elongator complex, is the enzyme that adds the initial cm^5^ group to affected tRNAs. Subsequent enzymes add other modifications, eventually leading to four final modifications (indicated) affecting 11 tRNA isoacceptors, and hence the translation of 22 possible codons[S1].

**Figure S4.**
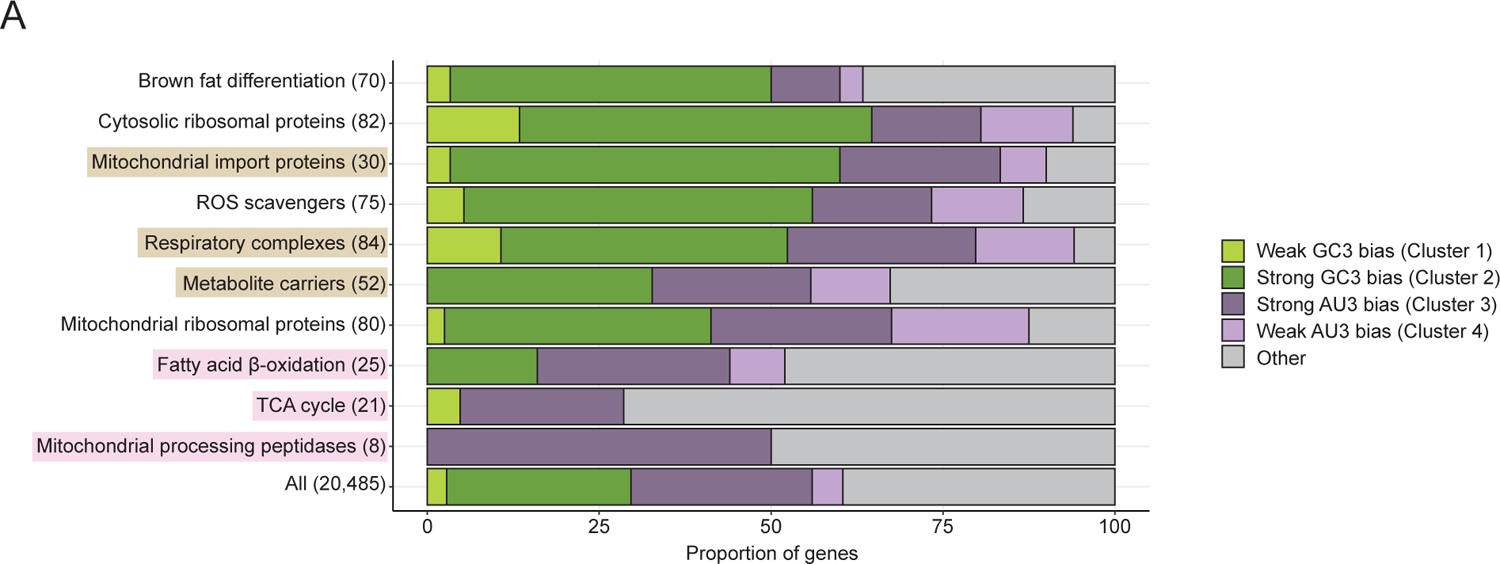
Codon usage differences in different functional gene groups, related to Figure 5. **(A)** Percentage bar plots showing differences in codon usage for different functional groups. For each functional group, gene distribution among different codon usage clusters are depicted. Clusters 1–4 are defined according to the classification in **Figure 5A**. All other genes that are not in these four clusters are grouped into the “Other” category. Mitochondrial membrane components (brown) and matrix components (pink) are highlighted.

**Figure S5.**
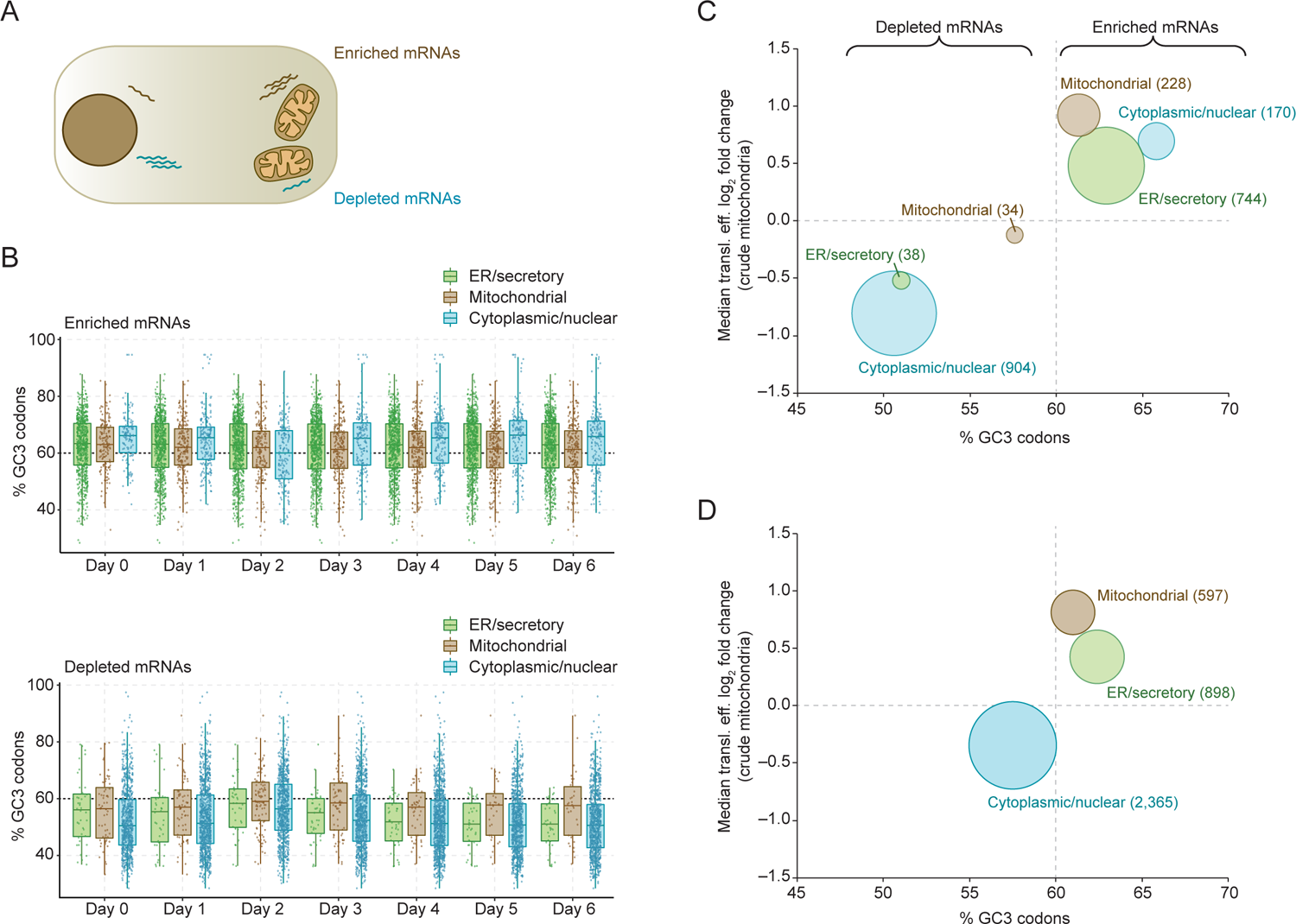
mRNAs encoding proteins that function in different cellular locations exhibit differences in mRNA partitioning and GC3 codon content, and are thus differentially regulated by the observed translation bias, related to Figure 6. (A) Schematic showing mRNAs that are enriched or depleted around the vicinity of mitochondria. (B) GC3codon content in mRNAs encoding proteins that function in different cellular locations. mRNAs are categorized as such: “ER/secretory”, all mRNAs encoding ER-resident components and components that would pass through the endomembrane system; “mitochondrial”, mRNAs encoding mitochondrial components; “cytoplasmic/nuclear”, all other mRNAs. Enriched mRNAs and depleted mRNAs are shown in separate plots. Otherwise, plotted as in Figure 6A. (C) Relationship between changes in translational efficiency in the crude mitochondria fraction and GC3 codon content, for enriched and depleted mRNAs. Enriched/depleted mRNAs are categorized as in (B). The size of the bubble indicates the number of genes in each category. (D) Relationship between changes in translational efficiency in the crude mitochondria fraction and GC3 codon content, for mRNAs encoding proteins that function in different cellular locations. Categories are as described in (B). Otherwise, as in (C).

---

**Table S1.**
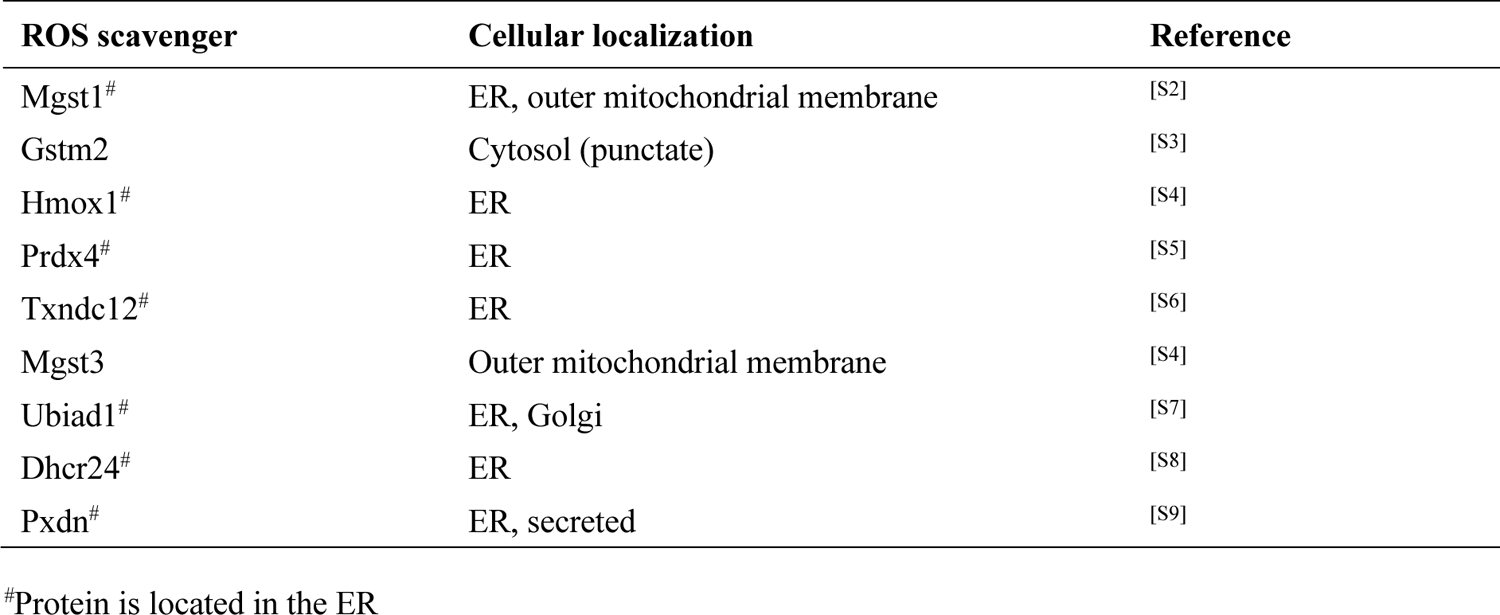
List of ROS scavengers encoded by mRNAs enriched in crude mitochondrial fraction.

